# Volume Electron Microscopy Analysis of Synapses in Associative and Primary Regions of the Human Cerebral Cortex

**DOI:** 10.1101/2024.03.26.586748

**Authors:** Nicolás Cano-Astorga, Sergio Plaza-Alonso, Javier DeFelipe, Lidia Alonso-Nanclares

## Abstract

Functional and structural studies investigating macroscopic connectivity in the human cerebral cortex suggest that high-order associative regions exhibit greater connectivity compared to primary ones. However, the synaptic organization of these brain regions remains unexplored due to the difficulties involved in studying the human brain at the nanoscopic level. In the present work, we conducted volume electron microscopy to investigate the synaptic organization of the human brain obtained at autopsy. Specifically, we examined layer III of Brodmann areas 17, 3b, and 4, as representative areas of primary visual, somatosensorial, and motor cortex. Additionally, we conducted comparative analyses with our previous synaptic datasets of layer III from temporopolar and anterior cingulate associative cortical regions (Brodmann areas 24, 38, and 21). 9,690 synaptic junctions were 3D reconstructed, showing that certain synaptic characteristics appeared to be specific to particular cortical regions. The number of synapses per volume, the proportion of the postsynaptic targets, and the synaptic size may distinguish one region from another, regardless of whether they are associative or primary cortex. By contrast, other synaptic characteristics were common to all analyzed regions, such as the proportion of excitatory and inhibitory synapses, their shapes, their spatial distribution, and a higher proportion of synapses located on dendritic spines. These observations may be included within the general rules of synaptic organization of the human cerebral cortex. The present results on nanoscopic characteristics of synapses provide further insights into the structural design of the human cerebral cortex.

**Significance statement:** Structural and functional studies investigating macroscopic connectivity in the human cerebral cortex have suggested that high-order associative regions exhibit greater connectivity compared to primary ones. However, the synaptic organization of these brain regions remains unexplored. Here, thousands of synaptic junctions were 3D reconstructed in associative and primary cortical regions. We found that certain synaptic characteristics appeared to be specific to particular cortical regions —regardless of whether they are associative or primary cortex— whereas others were common to all analyzed regions. The present results provide further insights into the structural design of the human cerebral cortex.

## Introduction

The study of the synaptic organization of the brain is far from over being complete. Of the different approaches to studying synapses at the ultrastructural level in the mammalian brain, the gold standard methodology is volume electron microscopy. However, this technology is very time-consuming and challenging to use for obtaining large volumes of brain tissue. Therefore, volume electron microscopy studies are usually applied to a relatively small number of sections, which increases statistical variability and affects the reliability of the results (reviewed in Merchán-Pérez et al., 2009). Nevertheless, automated or semi-automated electron microcopy techniques are a major advance in the study of synaptic organization, as series of consecutive sections comprising relatively large volume samples can be obtained (Denk and Horstmann 2004; Knott et al., 2008; Merchán-Pérez et al., 2009; Kleinfeld et al., 2011; Helmstaedter et al., 2013; Kasthuri et al., 2015; Titze and Genoud, 2016; Kubota et al., 2018; Rollenhagen et al., 2020). One strategy is to obtain dense reconstructions of brain tissue using serial block-face electron microscopy (SBEM) in a small region of an individual (e.g., Kasthuri et al., 2015; Motta et al., 2019; Karimi et al., 2020; Shapson-Coe et al., 2021; Loomba et al., 2022; Winding et al., 2023). Another strategy is to obtain multiple samples of smaller volumes of tissue using Focused Ion Beam/Scanning Electron Microscopy (FIB/SEM) in different regions of several individuals. This multiple sampling using volume electron microscopy is especially important when examining human brain tissue, not only because of the large size of the brain and extension of particular brain regions, but also because inter-individual variability is clearly more pronounced than in animal models. Thus, to explore the synaptic characteristics of a given region, our approach was to determine the range of variability by multiple sampling of relatively small volumes in any region of interest in several individuals.

Moreover, our current understanding of the brain structure stems from research conducted on experimental animals. However, certain fundamental structural and behavioral characteristics are unique to humans, and it is therefore imperative to acquire data directly from human brains (e.g., Oberheim et al., 2009; DeFelipe, 2015; Mansvelder et al., 2019). Functional and structural studies investigating macroscopic connectivity in the human cerebral cortex have suggested that high-order associative cortex exhibits greater connectivity compared to primary cortex (Sporns et al., 2005; Van Essen et al., 2013; Paquola et al., 2020). To obtain a more comprehensive understanding of brain organization, ideally the goal would be to integrate macro-, meso- and microscopic studies (DeFelipe, 2015). In this regard, autopsy samples are a suitable source of strictly normal tissue, but a delay in the post-mortem brain tissue fixation after death (5 or more hours) entails anatomical and metabolic changes (Gonzalez-Riano et al., 2017; 2021), making the tissue inappropriate for a feasible analysis. The scarcity of synaptic circuitry data in the normal human brain can be attributed to this primary factor. In addition, it has been previously shown that the 3D reconstruction method using FIB/SEM can be applied to study the synaptic organization of the human brain obtained at autopsy with short post-mortem delay in great detail, yielding good results (Domínguez-Álvaro et al., 2018, 2019, 2021a, 2021b; Montero-Crespo et al., 2020, 2021; Cano-Astorga et al., 2021, 2023).

In the present study, we used FIB/SEM to analyze non-pathological brain tissue samples from autopsy cases with a post-mortem delay of less than 4 h. We focused on Brodmann area (BA) 17, 3b, and 4 (Zilles and Amunts, 2010) as representative areas of primary visual (BA17), somatosensorial (BA3b) and motor cortex (BA4). These areas are involved in the processing of relatively basic sensory and motor tasks (reviewed in Mountcastle, 1997).

The first goal was to study the synaptic organization of the neuropil —where the vast majority of synapses are found (DeFelipe et al., 1999)— in these brain regions. The electron microscopic analyses were performed in layer III. The pyramidal cells located in layer III are the largest source of cortico-cortical axon projections (Felleman and Van Essen, 1991; Thomson and Lamy, 2007; Barbas, 2015; D’Souza and Burkhalter, 2017; Rockland, 2019) and this layer has been extensively studied using physiological and morphological analysis (e.g., see Eyal et al., 2018; Gidon et al., 2020; Galakhova et al., 2022; Benavides-Piccione et al., 2023 and references therein). Our second goal was to perform an integrative analysis including our previous layer III datasets from BA24, BA38 (ventral and dorsal) and BA21, which were studied in the same autopsy cases and using the same techniques (Cano-Astorga et al., 2023).

Hence, our purpose was to gain further insights into the ultrastructural synaptic characteristics of distinct cortical regions, aiming to better understand the synaptic organization of the human cerebral cortex.

## Material and Methods

### Tissue preparation

Human brain tissue was obtained from three autopsies (with short postmortem delays of less than 4 h) obtained from two men (53 and 66 years old) and one woman (53 years old) with no recorded neurological or psychiatric alterations. The procedure was approved by the Institutional Ethical Committee. Human brain tissue samples from the same autopsies had been used as control tissue in previous studies (Domínguez-Álvaro et al., 2018, 2019, 2021a; Montero-Crespo et al., 2020; Benavides-Piccione et al., 2020; 2023; Cano-Astorga et al., 2021, 2023; Plaza-Alonso et al., 2023).

Upon removal, brains were immediately immersed in cold 4% paraformaldehyde (Sigma-Aldrich, St Louis, MO, USA) in 0.1 M phosphate buffer (PB; Panreac, 131965, Spain), pH 7.4 for 24–48 h and sectioned into 1.5-cm-thick coronal slices.

Brain tissue samples from BA17 were taken from the depth of the calcarine fissure in the medial surface of the occipital pole (Kuljis, 1994). BA3b and BA4 were taken from the dorsal surface of the hemisphere and correspond to the caudal and rostral lips of the central sulcus, respectively (Jones, 1986). BA24 corresponds to the “p24b” field defined in Palomero-Gallagher et al. (2008), vBA38 and dBA38 correspond to the “TG” and “TAr” areas, respectively, defined in Ding et al., (2009), and BA21 corresponds to the middle temporal gyrus (T2).

Small blocks (10 × 10 × 10 mm) of each region of interest were then transferred to a second solution of 4% paraformaldehyde in PB for 24 h at 4°C. After fixation, the tissue blocks were washed in PB and sectioned coronally in a vibratome (Vibratome Sectioning System, VT1200S Vibratome, Leica Biosystems, Germany). 150 µm-thick sections were processed for electron microscopy, and 50 µm-thick sections were processed for Nissl staining to determine cytoarchitecture (Fig. 1).

**Figure 1.**
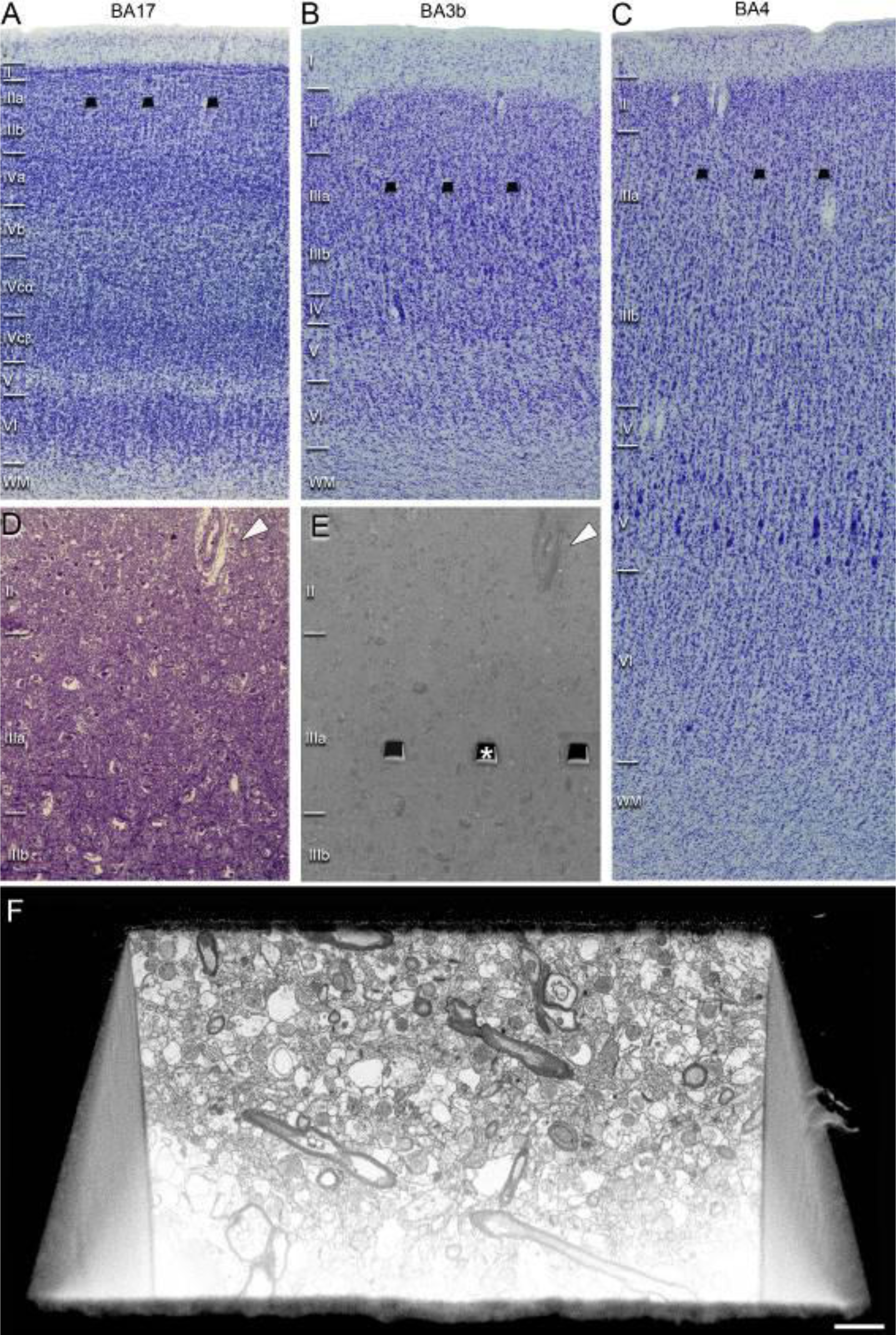
Cortical sampling regions. (A–C) Nissl-stained sections to illustrate the cytoarchitectonic differences between cortical regions BA17, BA3b and BA4 from an autopsy case (AB7). Layer delimitations are based on Kuljis (1994) for (A), and Jones (1986) for (B) and (C). The analyzed FIB/SEM sampling regions are shown as superimposed dark trapezoids in A–C. (D, E) Correlative light/electron microscopy analyses of the layer III neuropil. (D) 1-µm-thick semithin section stained with toluidine blue, which is adjacent to the block used for FIB/SEM imaging (E). (E) SEM image at higher magnification of (D) illustrating the block surface with trenches made in the neuropil to acquire the FIB/SEM stacks of images. White arrowheads in (D) and (E) point to the same blood vessel, allowing the exact location of the region of interest to be identified. (F) SEM image at higher magnification showing the front of a trench made to acquire an FIB/SEM stack of images. Scale bar (in F) indicates 300 µm in A–C, 250 µm in D and E, and 1.5 µm in F.

### Electron Microscopy Processing

Selected 150µm-thick sections were washed in 0.1M PB and postfixed for 48 h in a solution containing 2% paraformaldehyde, 0.2% glutaraldehyde (TAAB, G002, UK), and 0.003% CaCl_2_ (Sigma, C-2661-500G, Germany) in sodium cacodylate (Sigma, C0250-500G, Germany) buffer (0.1 M). Additionally, the sections were postfixed for 1 minute at 50°C and 150W power in a variable wattage microwave (PELCO BioWave Pro 36500-230) in a solution containing 2% paraformaldehyde, 2.5% glutaraldehyde, and 0.003% CaCl_2_ in sodium cacodylate buffer (0.1 M). The sections were treated with 1% OsO4 (Sigma, O5500, Germany), 0.1% potassium ferrocyanide (Probus, 23345, Spain), and 0.003% CaCl_2_ in sodium cacodylate buffer (0.1 M) for 1 h at room temperature. They were then stained with 1% uranyl acetate (EMS, 8473, USA), dehydrated, and flat-embedded in Araldite (TAAB, E021, UK) for 48 h at 60°C (DeFelipe and Fairén, 1993; Cano-Astorga et al., 2024). The embedded sections were then glued onto a blank Araldite block. Semithin sections (1–2 μm thick) were obtained from the surface of the block and stained with 1% toluidine blue (Merck, 115930, Germany) in 1% sodium borate (Panreac, 141644, Spain). The last semithin section (which corresponds to the section immediately adjacent to the block surface) was examined under light microscope and photographed to accurately locate the neuropil regions to be examined (Fig. 1).

### Volume fraction estimation of cortical elements

Twelve semithin sections (1–1.5 μm thick) from each case, stained with toluidine blue (see above), were used to estimate the volume fraction (Vv) occupied by neuropil, cell bodies (from neurons, glia and undetermined somata) and blood vessels. This estimation was performed applying the Cavalieri principle to 12 semithin sections per case (Gundersen et al., 1988) by point counting (Q) using the integrated Stereo Investigator stereological package (Version 8.0, MicroBrightField Inc., VT, USA) attached to an Olympus light microscope (Olympus, Bellerup, Denmark) at 40× magnification. A grid, whose points covered an area of 2500 μm^2^, was randomly placed at six sites over the traced layer III on six semithin sections (twelve semithin sections were obtained and every second one was analyzed) to determine the Vv occupied by the different elements: neuropil, cell bodies and blood vessels. Vv (in the case of the neuropil, for instance) was estimated with the following formula: Vv-neuropil = Q-neuropil *100 / (Q-neuropil + Q-neurons + Q-glia + Q-undetermined cells + Q-blood vessels).

### Three-Dimensional Electron Microscopy

The blocks containing the embedded tissue were glued onto a sample stub using conductive carbon tape (Electron Microscopy Sciences, USA #77825-09). All surfaces of the blocks except the top surface were covered with silver paint (Electron Microscopy Sciences, USA) to prevent any charging of the resin. The stubs with the mounted blocks were then placed into a sputter coater (Emitech K575X, Quorum Emitech, Ashford, Kent, UK) and the top surface was coated with several 10 nm-thick layers of gold/palladium to facilitate charge dissipation.

The 3D study of the samples was carried out using a dual beam microscope (Crossbeam® 40 electron microscope, Carl Zeiss NTS GmbH, Oberkochen, Germany). This instrument combines a high-resolution field-emission SEM column with a focused gallium ion beam (FIB), which permits removal of thin layers of material from the sample surface on a nanometer scale. Using a 7nA ion current (30 kV accelerating voltage), a first coarse trench was milled to provide visual access to the tissue below the block surface. The exposed surface of this trench was then milled with a 700 pA ion current (30 kV accelerating voltage) to remove 20 nm. As soon as one layer of material was removed by the FIB, the exposed surface of the sample was imaged by the SEM using the backscattered electron detector (at 1.6–1.7 kV acceleration potential, and 800– 1200 pA current probe). The sequential automated use of FIB milling and SEM imaging allowed us to obtain long series of microphotographs of a 3D sample of selected regions (Merchán-Pérez et al., 2009). Image resolution in the xy plane was 5 nm/pixel. Resolution in the z-axis (section thickness) was 20 nm, and image size was 2048×1536 pixels. These parameters were chosen in order to obtain a large enough field of view where synaptic junctions could be clearly identified, within a reasonable time frame (approximately 12 h per stack of images).

For the present study, a total of 22 series of images were acquired: 4 stacks in BA17 (approximately 600 μm from the pial surface from AB7; total volume studied: 1,763 μm^3^); 9 stacks in BA3b (approximately 600–700 μm from the pial surface, three stacks per case, AB2, AB3 and AB7; total volume studied: 4,034 μm^3^); and 9 stacks in BA4 (approximately 600–900 μm from the pial surface, three stacks per case, AB2, AB3 and AB7; total volume studied: 4,119 μm^3^). An example of the serial images is illustrated in Figure 2. The number of sections per stack ranged from 261 to 305 in BA17, 252–306 in BA3b, and 281–318 in BA4, which corresponds to a volume per stack ranging from 411 to 480 μm^3^ (mean: 441 μm^3^) in BA17, 396–481 μm^3^ (mean: 448 μm^3^) in BA3b, and 445–500 μm^3^ (mean: 458 μm^3^) in BA4.

**Figure 2.**
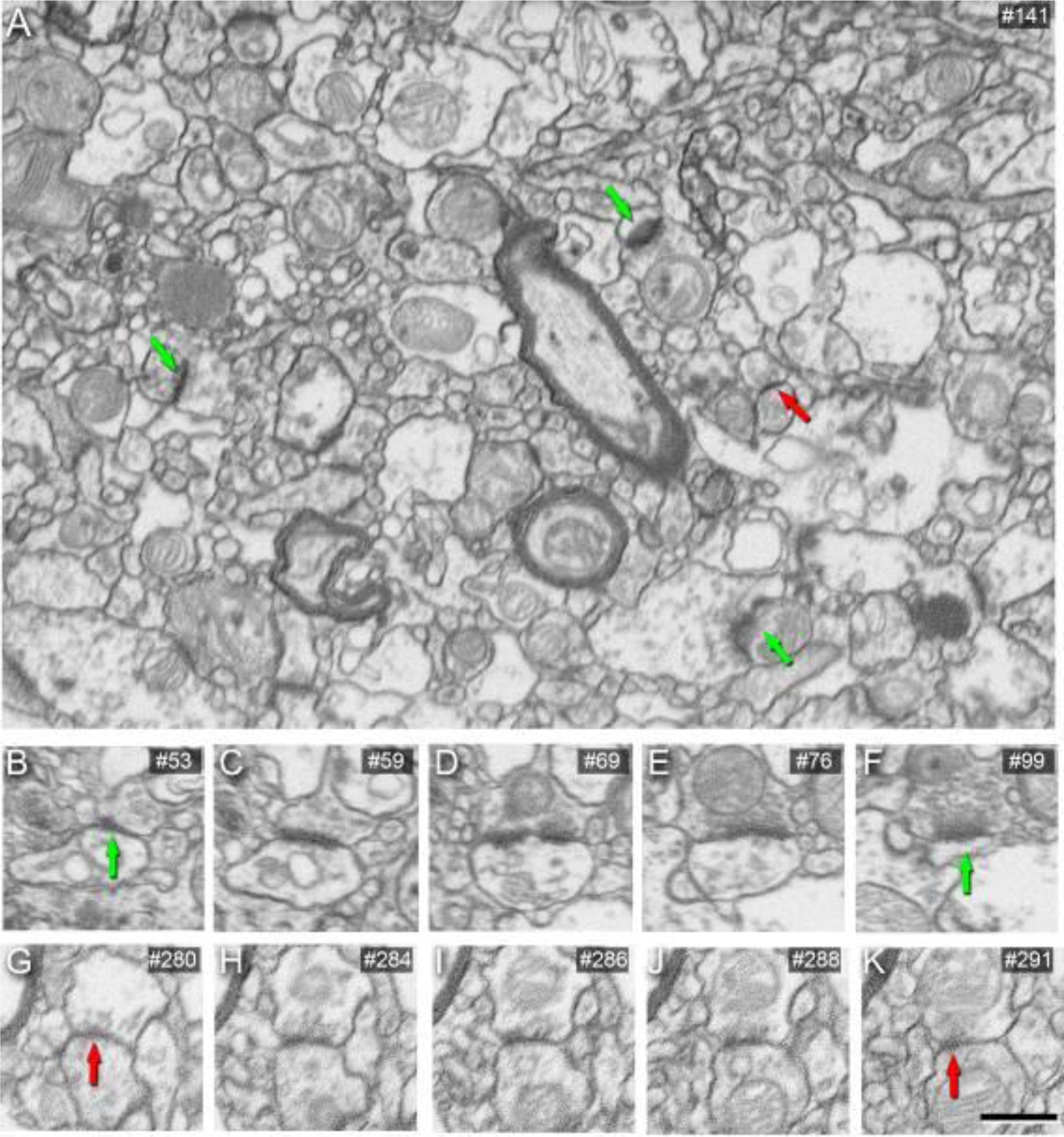
Images obtained by FIB/SEM showing AS and SS in the neuropil of human BA4. (A) Low-magnification FIB/SEM image from a stack of images. Green arrows point to some AS and the red arrow points out an SS. Sequence of FIB-SEM serial images of an AS (B–F) and an SS (G–K). Numbers on the top right of each panel indicate the number of each section from the stack of FIB/SEM images. Green arrows indicate the beginning (B) and the end (F) of the AS. Red arrows indicate the beginning (G) and the end (K) of the SS. Scale bar (in K) indicates 800 nm in A, and 600 nm in B– K.

Stacks of images from BA24, BA38v, BA38d, and BA21 (34 stacks) were previously acquired using the same technique, and similar volumes and numbers of sections were obtained (Cano-Astorga et al., 2023). All stack details are summarized in “Supplementary Table 1”.

Alignment (registration) of serial microphotographs obtained with the FIB/SEM was performed with the “Register Virtual Stack Slices” plug-in in FIJI, which is a version of ImageJ (ImageJ 1.51; NIH, USA) with a collection of preinstalled plug-ins (https://fiji.sc/). To avoid deformation of the original images, as well as size changes and other artifacts, we selected a registration technique that only allowed movement of individual images, with no rotation.

All measurements were corrected for tissue shrinkage, which occurs during the processing of sections (Merchán-Pérez et al., 2009). To estimate the shrinkage in our samples, we photographed and measured the area of the vibratome sections with FIJI, both before and after processing for electron microscopy. The section area values after processing were divided by the values before processing to obtain the volume, area, and linear shrinkage factors (Oorschot et al., 1991) — yielding correction factors of 0.90, 0.93, and 0.97, respectively. Nevertheless, in order to compare with previous studies — in which either no correction factors had been included or such factors were estimated using other methods— in the present study, we provided both sets of data. Additionally, a correction in the volume of the stack of images —to account for the presence of fixation artifact (e.g., swollen neuronal or glial processes)— was applied after quantification with Cavalieri principle (Gundersen et al., 1988; see Montero-Crespo et al., 2020). Every FIB/SEM stack was examined using FIJI, and the volume artifact was found to range from 0.0 to 16.1% of the volume stacks.

### Three-Dimensional Analysis of Synapses

The obtained FIB/SEM stacks of images were analyzed using EspINA software (EspINA Interactive Neuron Analyzer, 2.9.12; https://cajalbbp.csic.es/espina-2; Morales et al., 2011; Fig. 3).

**Figure 3.**
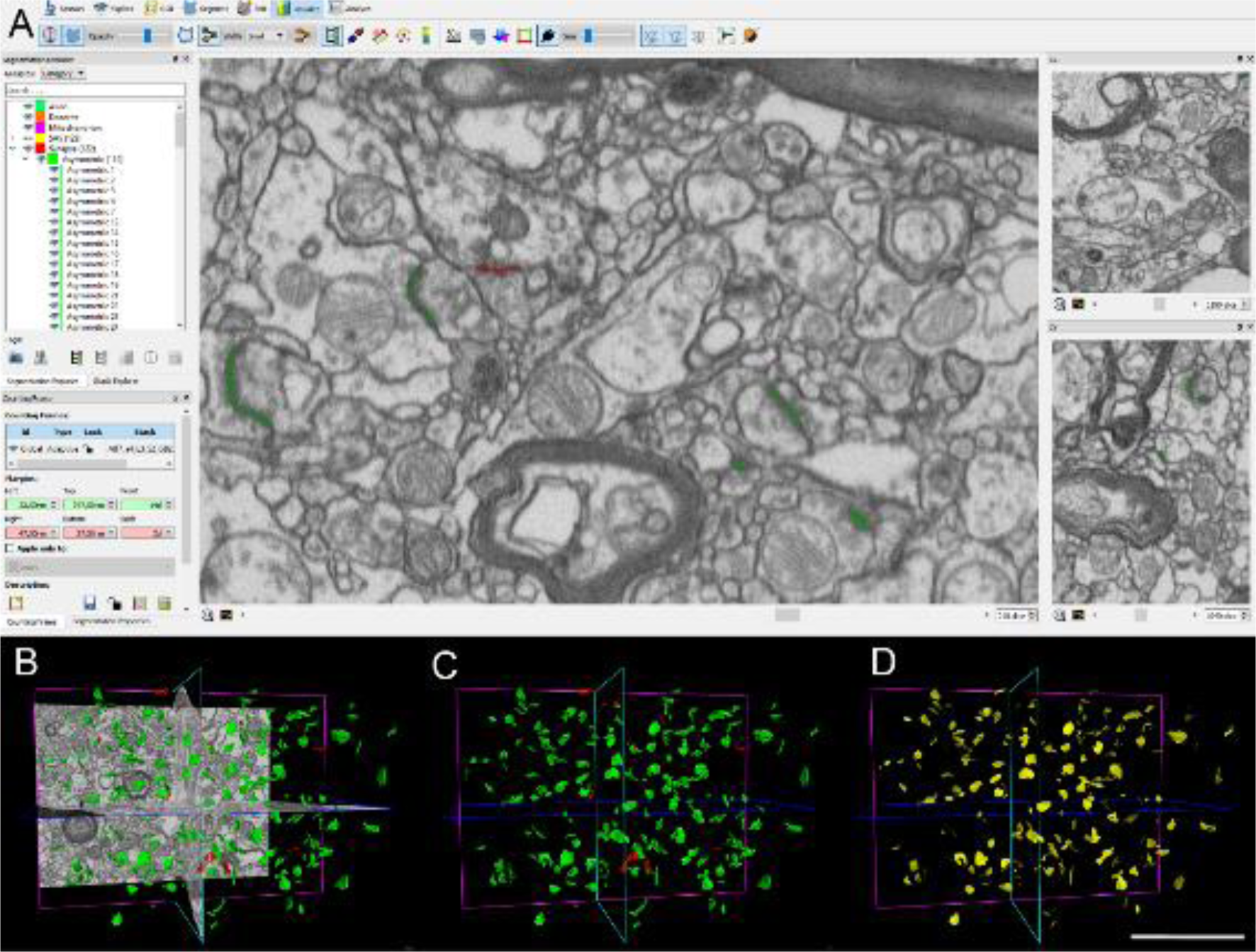
Three-dimensional analysis of synapses. (A–D) Screenshots of the EspINA software user interface. (A) In the main window, the sections are viewed through the xy plane (as obtained by FIB/SEM microscopy). The other two orthogonal planes, yz and xz, are also shown in windows on the right. (B) 3D window showing the three orthogonal planes and the 3D reconstructions of asymmetric (green) and symmetric (red) synaptic junctions. (C) 3D reconstructed synaptic junctions. (D) Computed synaptic apposition surface for each reconstructed synaptic junction (yellow). Scale bar (in D) indicates 5 μm for (B)–(D).

The user identifies synapses by assessing the presence of pre- and postsynaptic densities, along with the accumulation of synaptic vesicles in the presynaptic terminal. Synaptic segmentation depends on the intense electron density of these pre- and postsynaptic regions. Users must establish a grey-level threshold, and the segmentation algorithm subsequently selects pixels in the synaptic junction darker than the chosen threshold. The resulting segmentation creates a 3D object encompassing the active zone and postsynaptic densities, as these represent the two darkest areas of the synaptic junction. As previously discussed (Merchán-Pérez et al., 2009; Cano-Astorga et al., 2024), for classifying cortical synapses into asymmetric synapses (AS; or type I) and symmetric synapses (SS; or type II), the main characteristic was their prominent or thin postsynaptic density, respectively (Fig. 2). These two types of synapses correlate with different functions: AS are mostly glutamatergic and excitatory, while SS are mostly GABAergic and inhibitory (Colonnier, 1968; Gray, 1969; Peters and Kaiserman-Abramof, 1969; Houser et al., 1984; DeFelipe and Fariñas, 1992; Peters and Palay, 1996; Ascoli et al., 2008). Nevertheless, in single sections, the synaptic cleft and the pre- and postsynaptic densities are often blurred if the plane of the section does not pass at right angles to the synaptic junction. Since the software EspINA allows navigation through the stack of images, it was possible to unambiguously identify every synapse as AS or SS based on the thickness of the postsynaptic density (PSD) (Merchán-Pérez et al., 2009). Synapses with prominent PSDs are classified as AS, while those with thin PSDs are classified as SS (Gray, 1959; Colonnier, 1968; Peters et al., 1991; Fig. 2). All synapses were manually identified by an expert, and unclear synapses were reevaluated by the consensus of two or three experts.

In addition, geometrical features —such as size and shape— and spatial distribution features (centroids) of each reconstructed synaptic junction were also examined. The EspINA software tool facilitates the 3D reconstruction of the pre- and postsynaptic membranes of the synaptic junction, which we will refer to as “3D reconstructed synaptic junction”. This software also extracts the Synaptic Apposition Area (SAS) and provides its measurements. Given that the pre- and postsynaptic densities are located face to face, their surface areas are comparable (for details, see Morales et al., 2013; Fig. 3D). Since the SAS comprises both the active zone and the PSD, it is a functionally relevant measure of the size of a synaptic junction (Morales et al., 2013). EspINA was also used to visualize each of the reconstructed synaptic junctions in 3D and to detect the possible presence of perforations or deep indentations in their perimeters. Regarding the shape of the PSD, the synapses could be classified into four main categories, according to the categories proposed by Santuy et al. (2018a): macular (disk-shaped PSD), perforated (with one or more holes in the PSD), horseshoe-shaped (with an indentation), and fragmented (disk-shaped PSDs with no connection between them). To identify the postsynaptic targets of the synapses, we navigated through the image stack using EspINA to determine whether the postsynaptic element was a dendritic spine (spine, for simplicity) or a dendritic shaft. As previously described in Domínguez-Álvaro et al. (2021a, 2021b), unambiguous identification of spines requires the spine to be visually traced to the parent dendrite (see Cano-Astorga et al., 2021), in which case we refer to them as complete spines. When synapses were established on a spine head-shaped postsynaptic element whose neck could not be followed to the parent dendrite, we identified these elements as incomplete spines. These incomplete spines were identified on the basis of their size and shape, the lack of mitochondria, and the presence of a spine apparatus — or because they were filled with ‘fluffy material’ (a term coined by Peters et al. (1991) to describe the fine and indistinct filaments present in the spines; see also del Río and DeFelipe, 1995). For simplicity, we will refer to both the complete and incomplete spines as spines, unless otherwise specified. We also recorded the presence of single or multiple synapses on a single spine. Furthermore, we determined whether the target dendrite had spines or not.

### Quantification of the Synaptic Density

EspINA provided the 3D reconstruction of every synaptic junction and allowed the application of an unbiased 3D counting frame (CF) — a regular prism enclosed by three acceptance planes and three exclusion planes marking its boundaries. All objects within the CF are counted, as are those intersecting any of the acceptance planes, while objects that are outside the CF, or intersecting any of the exclusion planes, are not counted. Thus, the number of synapses per unit volume was calculated directly by dividing the total number of synapses counted by the volume of the CF (Merchán-Pérez et al., 2009). This method was used in all 22 stacks of images.

### Spatial Distribution Analysis of Synapses

To analyze the spatial distribution of synapses, spatial point pattern analysis was performed as described elsewhere (Antón-Sánchez et al., 2014; Merchán-Pérez et al., 2014). Briefly, we compared the actual position of centroids of synaptic junctions with the CSR model — a random spatial distribution model that defines a situation where a point is equally likely to occur at any location within a given volume. For each of the 22 FIB/SEM stacks of images, we calculated three functions commonly used for spatial point pattern analysis: G, F, and K functions (Supplementary Fig. 1). As described in Merchán-Pérez et al. (2014) (see also Antón-Sánchez et al., 2014), the G function, also called the nearest-neighbor distance cumulative distribution function or the event-to-event distribution, is —for a distance d— the probability that a typical point separates from its nearest neighbor by a distance of d at the most. The F function, also known as the empty space function or the point-to-event distribution, is —for a distance d— the probability that the distance of each point (in a regularly spaced grid of L points superimposed over the sample) to its nearest synaptic junction centroid is d at the most.

The K function, also called the reduced second moment function or Ripley’s function, is—for a distance d— the expected number of points within a distance d of a typical point of the process divided by the intensity λ. An estimation of the K function is given by the mean number of points within a sphere of increasing radius d centered on each sample point, divided by an estimation of the expected number of points per unit volume. This study was carried out using the Spatstat package and R Project program (Baddeley et al., 2015).

### Statistical Analysis

Kruskal–Wallis (KW) nonparametric test, with post-hoc correction (via Dunn’s multiple comparisons) was performed to compare —between regions— the synaptic density and the SAS area. Mann-Whitney (MW) nonparametric tests were performed to compare the SAS area between AS and SS; and SAS area of macular and complex-shaped synapses. To perform statistical comparisons of AS:SS proportions regarding their synaptic shape and their postsynaptic targets, chi-square (χ^2^) test was used for contingency tables. The same method was used to compare —between regions— the volume fraction occupied by the cortical structures, AS:SS proportion, the proportion of different synaptic shapes, and the proportion of postsynaptic targets. Frequency distribution analysis of the SAS area comparison studies were performed using Kolmogorov–Smirnov (KS) nonparametric test.

The above-mentioned statistical analyses were performed with the GraphPad Prism statistical package (Prism 9.00 for Windows, GraphPad Software Inc., USA), as well as the on-line tool VassarStats (http://vassarstats.net/).

Data variability between individuals was estimated by calculating the coefficient of variation (CV) of each synaptic parameter in every cortical region (except BA17, which only had data from one case). CV was calculated by dividing the standard deviation (SD) by the mean for each of the synaptic parameters analyzed (Table 2).

Goodness-of-fit tests were performed to find the theoretical probability density functions that best fitted the empirical distributions of SAS areas in all cortical regions. These analyses were performed with EasyFit Professional 5.5 (MathWave Technologies).

## Results

The following is a detailed description of the characteristics of the neuropil and the synapses of layer III from BA17, BA3b, and BA4, as well as a comparative analysis with previously studied temporopolar and cingulate regions (BA24, vBA38, dBA38, and BA21; Cano-Astorga et al., 2023).

### Volume fraction of cortical elements

Since the neuropil is where the vast majority of synapses are found (DeFelipe et al., 1999), the volume fraction (Vv) was estimated to determine the relative volume occupied by different cortical elements: neuropil, cell bodies (including neurons, glial cells and undetermined cells), and blood vessels. The neuropil constituted the main component (ranging from 73.53% in BA17 to 79.94% in BA4; Supplementary Table 2), followed by cell bodies (ranging from 14.82% in BA4 to 18.13% in BA3b), and blood vessels (ranging from 4.22% in BA3b to 8.91% in BA17; Supplementary Table 2).

To determine whether there was a difference in the Vv occupied by the cortical elements, data from BA17, BA3b and BA4 were compared with our previous studies from BA24, vBA38, dBA38 and BA21 performed in the same individuals (Cano-Astorga et al., 2023). Contingency tables were applied and statistical differences were found (χ^2^, p<0.0001) indicating that the Vv of neuropil was larger in vBA38, dBA38 and BA21 than in BA24, BA17, BA3b and BA4 (Fig. 4A).

**Figure 4.**
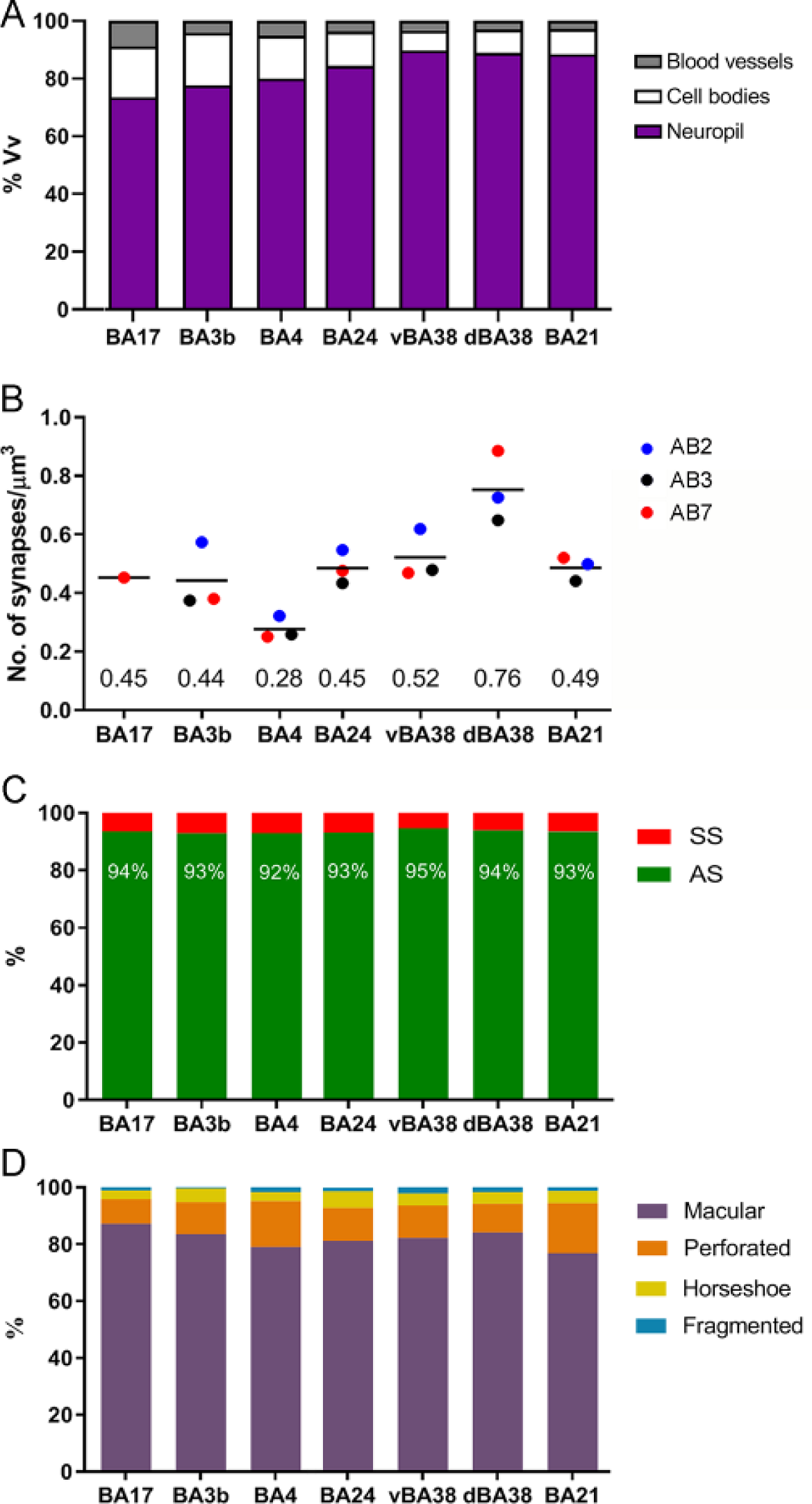
Comparison of the synaptic characteristics of the layer IIIA neuropil in different cortical regions. (A) Plot of the volume fraction occupied by each cortical element. The volume fraction occupied by neuropil was larger in vBA38, dBA38, and BA21 than in the other cortical regions (χ^2^, p<0.0001). (B) Mean synaptic density of the neuropil. Each dot represents a single autopsy case according to the color key on the right. BA4 and dBA38 had the lowest and highest synaptic density, respectively (KW; P<0.05). (C) Proportions of AS and SS showed no differences between cortical regions (χ^2^; P>0.05). (D) Study of the synaptic shape of AS (macular, perforated, horseshoe-shaped, or fragmented) across cortical regions. BA4 and BA21 had a higher proportion of complex-shaped synapses (including perforated, horseshoe, and fragmented synapses) than the other regions (χ^2^; P<0.05).

### Synaptic density

Synapses were studied in 22 stacks of images from the layer III neuropil (i.e., excluding cell bodies, blood vessels, and major dendritic trunks). A total of 4,046 synaptic junctions were identified and 3D reconstructed from BA17 (900), BA3b (1900) and BA4 (1246). Of these, 693 (BA17), 1,424 (BA3b), and 878 (BA4) synapses were analyzed after discarding incomplete synapses or those touching the exclusion edges of the counting frame (see above) (Table 1; Supplementary Table 3).

**Table 1.** Accumulated data obtained from the ultrastructural analysis of neuropil from layer III of BA17, BA3b, and BA4 in human autopsy samples from three individuals. Data in parentheses have not been corrected for shrinkage. BA17 data come from the analyses of three stacks of images in one individual. The data for individual cases are shown in Supplementary Table 3. AS: asymmetric synapses; BA: Brodmann area; CF: counting frame; SAS: synaptic apposition surface; SD: standard deviation; SE: standard error of the mean; SS: symmetric synapses.

| **Region** | No. AS | No. SS | No.<br>synapses<br>(AS+SS) | % AS | % SS | CF<br>volume<br>($\mu\text{m}^3$) | No. AS/$\mu\text{m}^3$<br>(mean ± SD) | No. SS/$\mu\text{m}^3$<br>(mean ± SD) | No. all<br>synapses/$\mu\text{m}^3$<br>(mean ± SD) | Intersynaptic<br>distance (nm;<br>mean ± SD) | Area of SAS AS<br>($\text{nm}^2$; mean ± SE) | Area of SAS SS<br>($\text{nm}^2$; mean ± SE) |
| --- | --- | --- | --- | --- | --- | --- | --- | --- | --- | --- | --- | --- |
| **BA17** | 649 | 44 | 693 | 93.66% | 6.34% | 1,521<br>(1370) | $0.43 \pm 0.06$<br>($0.47 \pm 0.07$) | $0.03 \pm 0.01$<br>($0.03 \pm 0.01$) | $0.45 \pm 0.07$<br>($0.47 \pm 0.07$) | $843 \pm 50$<br>($784 \pm 47$) | $98,932 \pm 2125$<br>($92,007 \pm 1,976$) | $69,781 \pm 6,617$<br>($64,896 \pm 6,154$) |
| **BA3b** | 1328 | 96 | 1424 | 92.87% | 7.13% | 3,221<br>(2,900) | $0.41 \pm 0.11$<br>($0.46 \pm 0.13$) | $0.03 \pm 0.00$<br>($0.03 \pm 0.00$) | $0.44 \pm 0.11$<br>($0.49 \pm 0.13$) | $864 \pm 53$<br>($803 \pm 50$) | $99,224 \pm 1,573$<br>($92,278 \pm 1,463$) | $63,561 \pm 4,823$<br>($59,112 \pm 4,485$) |
| **BA4** | 809 | 69 | 878 | 92.09% | 7.91% | 3,199<br>(2,881) | $0.26 \pm 0.04$<br>($0.28 \pm 0.04$) | $0.02 \pm 0.00$<br>($0.02 \pm 0.00$) | $0.28 \pm 0.04$<br>($0.31 \pm 0.04$) | $946 \pm 61$<br>($879 \pm 57$) | $130,869 \pm 3082$<br>($121,708 \pm 2,866$) | $75,560 \pm 5,529$<br>($121,708 \pm 2,866$) |

The synaptic density analyses revealed no differences in the values for the number of synapses per volume in the analyzed regions: 0.45 synapses/μm^3^ in BA17, 0.38–0.57 synapses/μm^3^ in BA3b, and 0.25–0.32 synapses/μm^3^ in BA4 (Table 1; Supplementary Table 3).

Data from BA17, BA3b and BA4 were compared with our previous datasets from BA24, vBA38, dBA38, and BA21 pertaining to the same cases (Cano-Astorga et al., 2023). Statistical analysis showed that BA4 (0.28 synapses/μm^3^) and dBA38 (0.76 synapses/μm^3^) had the lowest and highest synaptic density, respectively (KW; P<0.05; Fig. 4B). No significant differences were found in the mean synaptic density between BA17, BA3b, BA24, vBA38, and BA21 (KW; P>0.05) (Fig. 4B).

### Proportion of AS and SS

The proportion of AS:SS was 92:2 in BA4, and 93:7 in BA17 and BA3b (Table 1; Supplementary Table 3). No significant differences were found in the AS:SS proportion between any cortical region, including BA24, vBA38, dBA38, and BA21 (Cano-Astorga et al., 2023) (χ^2^; P>0.05; Fig. 4C).

### Three-dimensional spatial distribution of synapses

To analyze the spatial distribution of the synapses, the actual position of each of the synaptic junctions in each stack of images was compared with the Complete Spatial Randomness (CSR) model. For this, the functions G, K, and F were calculated in the 22 stacks of images from BA17, BA3b, and BA4. We found that 2 stacks of images from BA4 did not fit into the CSR model, showing a slight tendency for a regular pattern in one of the functions (Supplementary Fig. 1A). In the remaining samples (i.e., 20 out of 22 stacks), the 3 spatial statistical functions resembled the theoretical curve that simulates the random spatial distribution pattern, which indicated that synapses fitted a random spatial distribution model in BA17, BA3b, and BA4 (Supplementary Fig. 1B).

In addition, the mean distance of each synaptic junction centroid to its nearest neighboring synaptic junction was further compared between all cortical regions. The comparative analysis with the rest of the cortical regions showed that BA4 displayed the largest distance (946 nm) — although statistical differences were only found compared to dBA38, which displayed the smallest distance (739 nm; Cano-Astorga et al., 2023) (KW; P<0.05; Table 1; Supplementary Table 3). No significant differences were found between the remaining cortical areas.

### Synaptic shape

To analyze the shape of the synaptic contacts, we classified each identified synapse into four categories: macular (with a flat, disk-shaped postsynaptic density [PSD], perforated (with one or more holes in the PSD), horseshoe (with an indentation in the perimeter of the PSD), or fragmented (with two or more physically discontinuous PSDs; for a detailed description, see Santuy et al., 2018a; Domínguez-Álvaro et al., 2019). The analyses showed that the vast majority of AS had a macular shape in all of the cortical regions analyzed (range: 79–87%), followed by perforated (range: 9–16%), horseshoe (range: 3–5%) and fragmented (range: 0.4–1.6%; Supplementary Table 4). Regarding the SS, the results were very similar in all analyzed regions: the majority of SS had a macular shape (range: 80–88%; Supplementary Table 4), whereas the perforated and horseshoe shapes were less frequent, with a range of 6–7% and 3–11%, respectively. SS with a fragmented shape were scarce; indeed, this shape was only identified in 6 out of 196 identified SS (Supplementary Table 4).

To determine whether there was a difference in the proportion of synaptic shapes between regions, contingency tests were applied comparing present and previous results from Cano-Astorga et al. (2023). Perforated, horseshoe, and fragmented synapses were computed as a group (as complex-shaped synapses) and we found that complex-shaped AS were more frequent in BA4 (21%) and BA21 (22.4%) than in the rest of the cortical regions (χ^2^; P<0.05; Fig. 4D). No significant differences were found in the proportion of complex-shaped SS (range: 8.3–20.5%) between cortical regions (χ^2^; P>0.05).

### Study of the postsynaptic targets

Postsynaptic targets were identified and classified as spines (corresponding to axospinous synapses) or dendritic shafts (axodendritic synapses). We also determined whether a synapse was located on the neck or head of a spine (Fig. 5). When the postsynaptic element was identified as a dendritic shaft, it was sub-classified as “with spines” or “without spines” (Supplementary Table 5).

**Figure 5.**
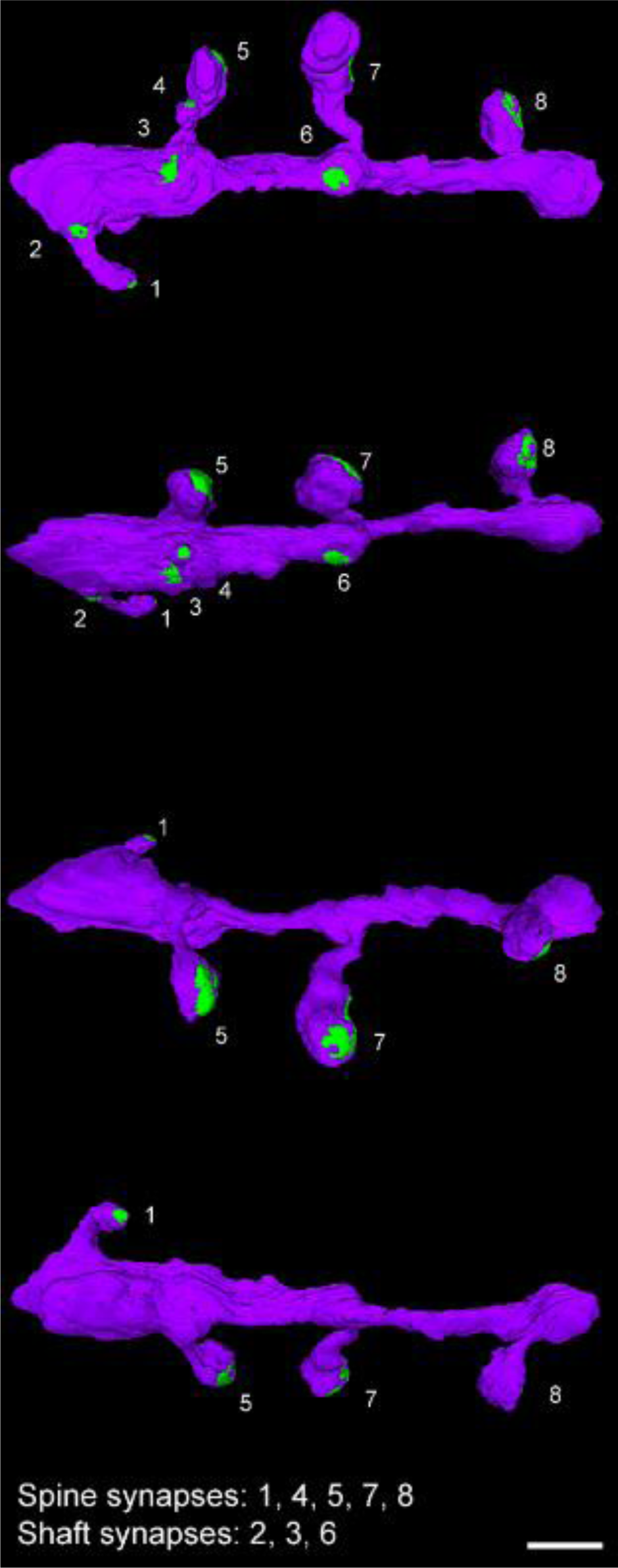
Example of a 3D reconstructed dendritic segment (purple) establishing asymmetric (green) synapses. No symmetric synapses were found in this dendritic segment. Synaptic junctions of the dendritic segment are shown from different angles. Numbers correspond to the same synapse. Scale bar (in C): 1 µm.

The postsynaptic elements of 637, 1,322, and 834 synapses were determined in BA17, BA3b, and BA4, respectively. Most synapses were AS established on dendritic spine heads (range: 57.1–68.5%) followed by AS on dendritic shafts (range: 24.2–33.9%), SS on dendritic shafts (range: 4–6.1%), and SS established on dendritic spine heads (range: 1.1–2.2%; Fig. 6). The least frequent types of synapses were AS on dendritic spine necks (range: 0.2–0.8%) and SS on dendritic spine necks (range: 0–0.5%; Fig. 6).

**Figure 6.**
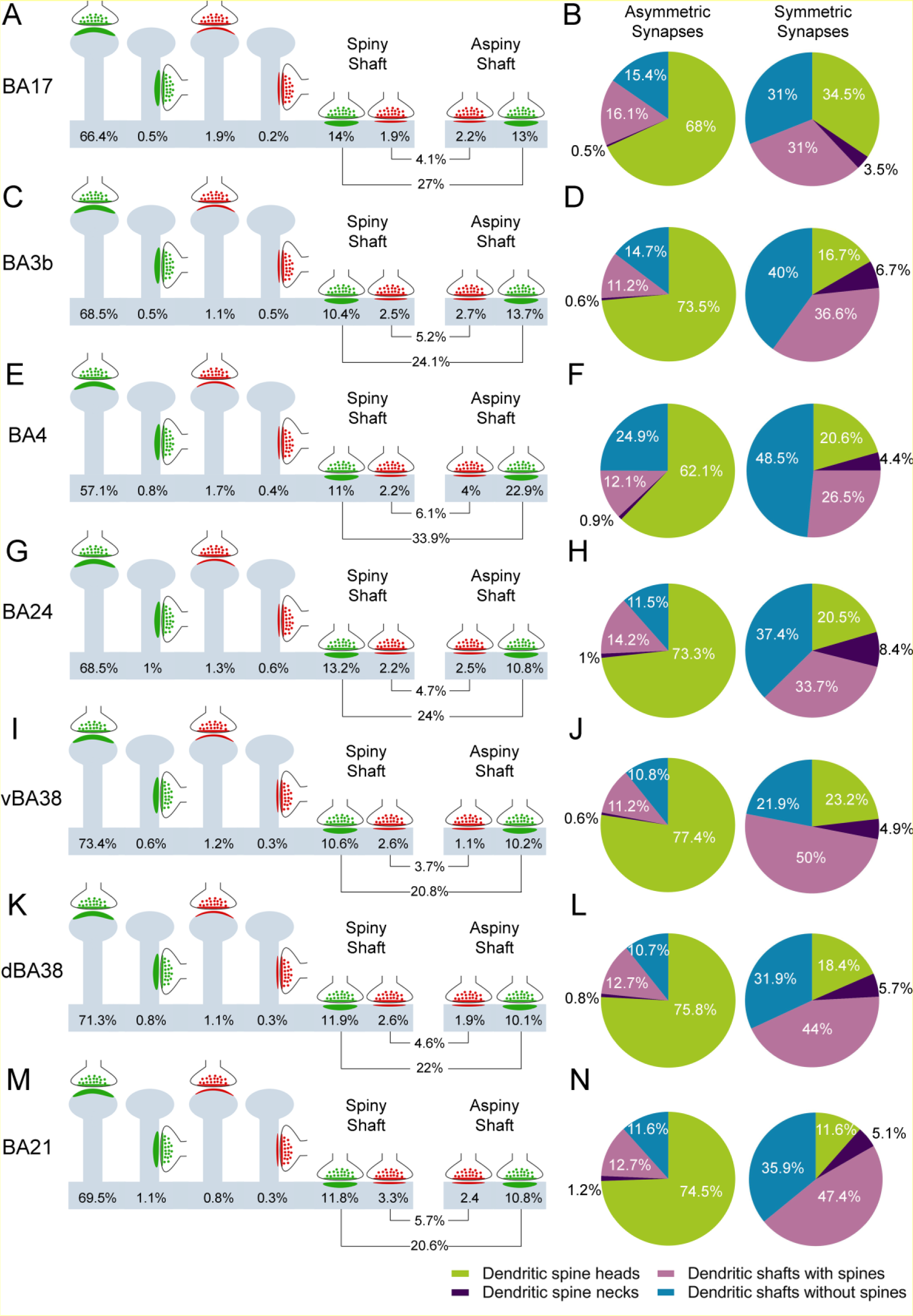
Representation of the distribution of synapses according to their postsynaptic target in different human cortical regions. (A, C, E, G, I, K, M) Data refer to the percentages of axospinous (i.e., head and neck of spines) and axodendritic (spiny and aspiny shafts) asymmetric (green) and symmetric (red) synapses. (B, D, F, H, J, L, N) Pie charts to illustrate the proportions of AS and SS according to their location as axospinous synapses (i.e., on the head or neck of the spine) or axodendritic synapses (i.e., spiny or aspiny shafts). AS established on dendritic shafts without spines were more frequent in BA4 and BA17 than in the other cortical regions (χ^2^, P<0.05). Data from BA24, vBA38, dBA38, and BA21 taken from Cano-Astorga et al., 2023. BA: Brodmann area; d: dorsal; Vv: volume fraction; v: ventral.

Proportions of the postsynaptic targets were compared including the previously analyzed regions BA24, vBA38, dBA38, and BA21 (Cano-Astorga et al., 2023). AS established on dendritic shafts without spines were more frequent in BA4 and BA17 than in the other cortical regions (χ^2^, P<0.05; Fig. 6). AS established on spine necks were less frequent in vBA38 and dBA38 than in the other cortical regions (χ^2^, P<0.05; Fig. 6). Concerning SS, those established on dendritic shafts without spines were more frequent in BA4 than in the other cortical regions (χ^2^, P<0.05), whereas SS established on dendritic shafts with spines were more frequent in vBA38 (χ^2^, P<0.05).

The spines were further analyzed to determine the number and type of synapses established on them, aiming to estimate the number of multiple synapses. The vast majority of spines had a single AS (BA4: 89.6%; BA17: 92.2%; BA3b: 95.2%), followed by spines with a large variety in the synapse number and location (head or neck) of AS and SS (Fig. 7). Comparisons of the multisynaptic spine proportions between cortical regions showed that spines with more than one AS (Fig. 7) were significantly more frequent in BA4 (10.4%) than in the rest of the regions (χ^2^, P<0.01).

**Figure 7.**
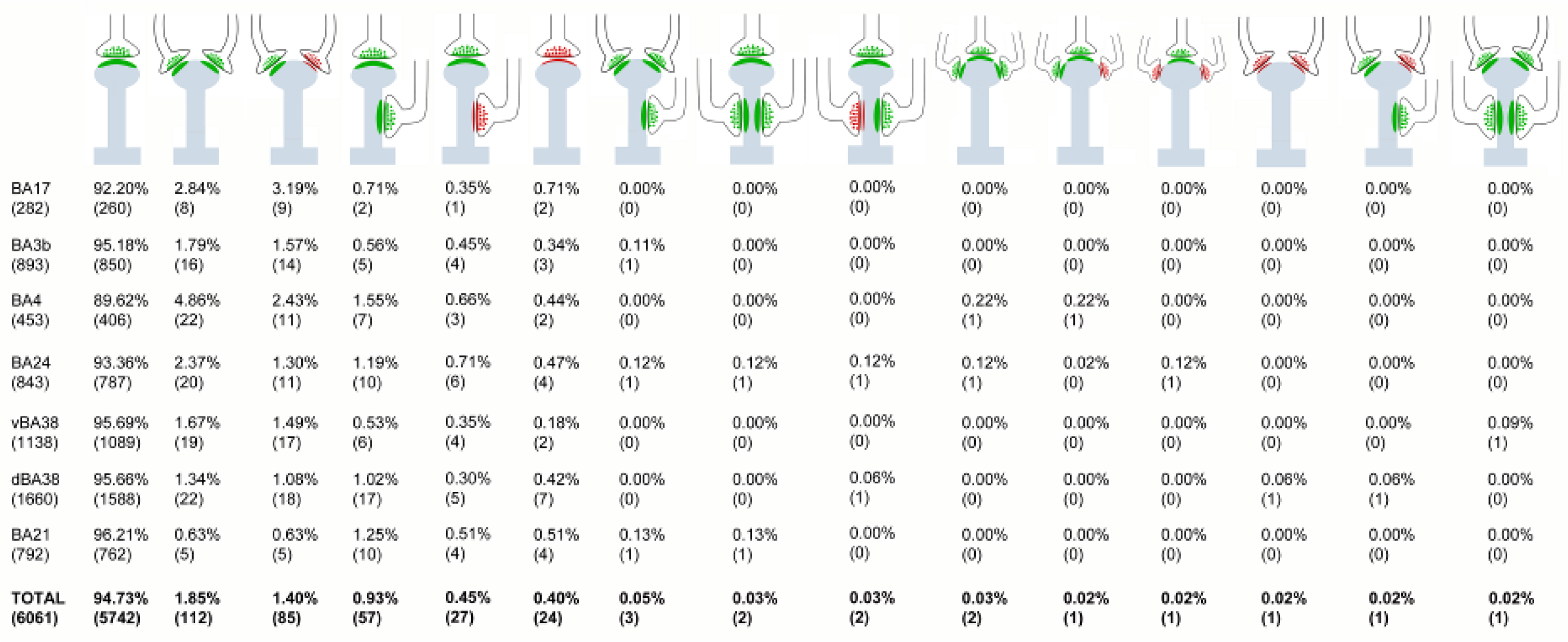
Study of the Multisynaptic Spines. Schematic representation of dendritic spine heads (including both complete and incomplete spines) receiving single and multiple synapses in BA17, BA3b, BA4, BA24, vBA38, dBA38 and BA21. Percentages of each type are indicated (absolute numbers of synapses are in parentheses). AS have been represented in green and SS in red. Spines with more than one AS (including all combinations) were more frequent in BA4 than in the other regions (χ^2^, P<0.05). AS: asymmetric synapses; BA: Brodmann area; d: dorsal; SS: symmetric synapses; v: ventral.

### Synaptic size

The mean size of the synapses (measured by the SAS area) showed that AS were larger than SS in BA17, BA3b, and BA4 (MW; P<0.001; Table 1; Supplementary Table 2). Comparison of the frequency distribution between AS and SS showed that larger synaptic junctions were also more frequent in AS than SS in BA17, BA3b, and BA4 (KS; P<0.01; Supplementary Fig. 2). To characterize the data distribution of AS and SS SAS area, we performed goodness-of-fit tests to find the theoretical probability density functions that best fitted the empirical distributions of SAS areas in all regions. We found that the best fit corresponded to a log-normal distribution, with some variations in the location (µ; range: 11.346–11.539 in AS and 10.757–11.058 in SS) and scale (σ; range: 0.5626–0.7533 in AS and 0.5999–0.9529 in SS) parameters. This was the case in all regions for both AS and SS, although the fit was better for AS than for SS, probably due to the smaller number of SS.

The frequency distribution of the SAS areas of AS in BA17, BA3b and BA4 were then compared with our previous datasets from BA24, vBA38, dBA38, and BA21 (Cano-Astorga et al., 2023). Larger AS were more frequent in BA4 than in any other cortical region, whereas smaller AS were more frequent in dBA38 (Fig. 8A, KS; P<0.0001). Smaller AS were more frequent in BA17 and BA3b than in BA24, vBA38 and BA21 (KS; P<0.0001; Fig. 8A). Similarly, significant differences (KW; P<0.01) were found between the mean SAS area of AS, indicating that the largest AS were in BA4, and the smallest were in dBA38. Also, AS in vBA38 were smaller than in BA24 (KW; P<0.01). The SAS area of SS was similar in all regions, and no differences were found in either their frequency distribution (KS; P>0.0001; Fig. 8B) or their mean SAS area (KW; P>0.0001).

**Figure 8.**
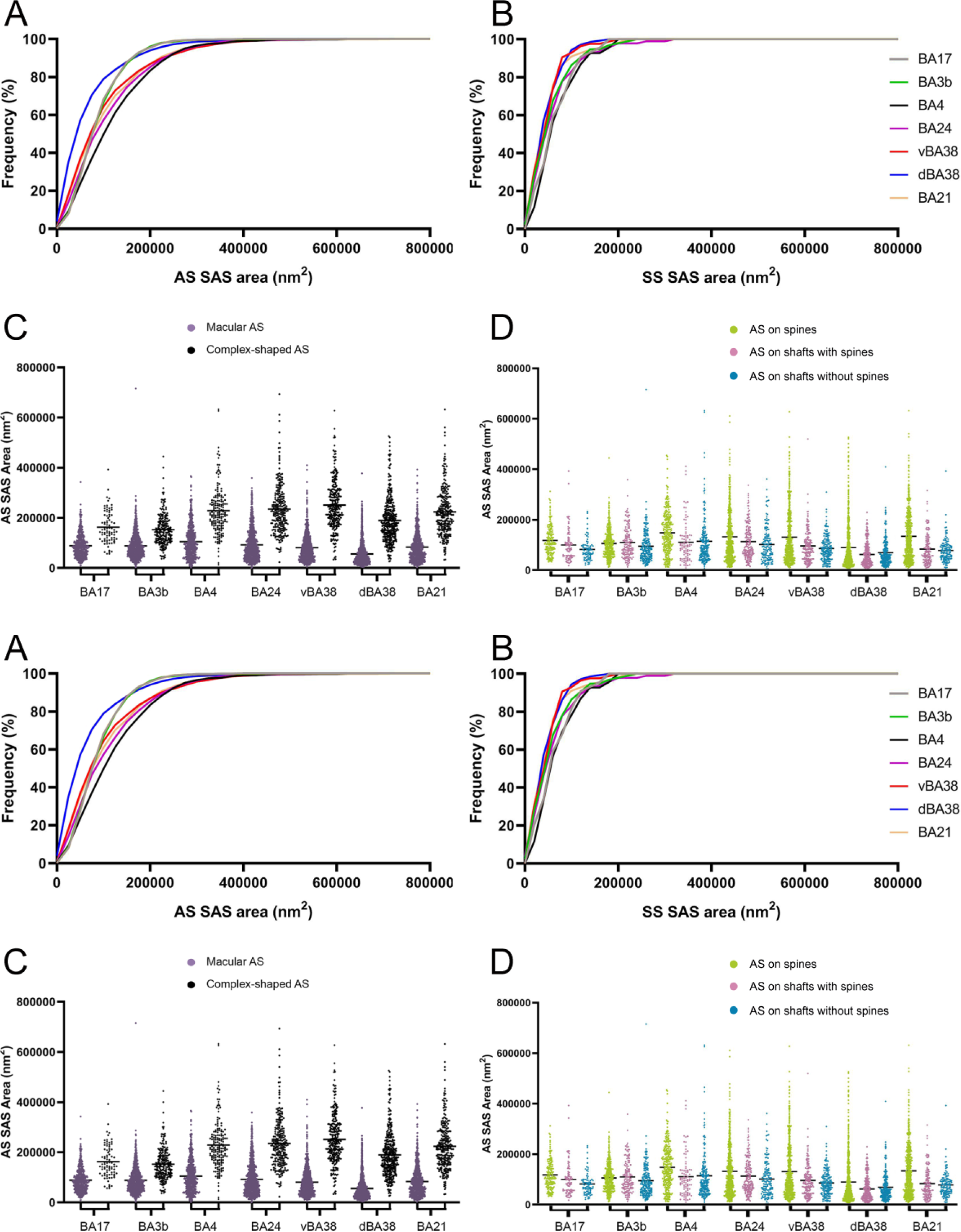
Comparison of the size of synapses in the neuropil of layer IIIA from different human cortical regions. (A) Cumulative frequency distribution of SAS area of AS according to the colored key in (B). The larger AS and smaller AS were more frequent in BA4 and dBA38, respectively (KS; P<0.0001). (B) Cumulative frequency distribution of SAS area of SS according to the colored key on the right. The frequency distributions of SAS area of SS were similar in all cortical regions and no differences were found. (C) Area of SAS of AS by morphology (macular and complex-shaped, see the colored key at the top). The area of the macular AS was significantly smaller than the area of the complex-shaped AS in all cortical regions (KW; P<0.001). (D) SAS Area of AS according to their postsynaptic target (spine heads, dendritic shafts with spines and without spines; refer to the colored key at the top). The area of the AS established on spine heads was larger than the area of the AS on dendritic shafts in all regions (KW; P<0.001), except in BA3b and BA24.

### Synaptic size and shape

We also determined whether the synaptic size is related to its shape. We found that the mean SAS area of the macular AS was significantly smaller than the complex-shaped AS in BA17, BA3b and BA4 (MW; P<0.001; Figure 8C; Supplementary Table 6). Similarly, the smaller AS were more frequent in the macular-shaped synapses than in the complex-shaped AS in BA17, BA3b, and BA4 (KS; P<0.01; Supplementary Fig. 3; Supplementary Table 6). Since the number of complex-shaped SS was very low (ranging from 9 to 14 in the cortical regions analyzed), this analysis was discarded as it would not have been sufficiently robust.

We also re-analyzed data from BA24, vBA38, dBA38, and BA21, comparing both *between* these four areas and with the present results. The mean SAS area of macular AS ranged from a minimum of 56,222 nm^2^ in dBA38 to a maximum of 104,855 nm^2^ in BA4. In the case of complex-shaped AS, the mean SAS area ranged from 159,924 nm^2^ in BA3b to 235,605 nm^2^ in BA24. We found that the mean SAS area of complex-shaped AS was smaller in BA3b and BA17 than in the other cortical regions (KW; P<0.0001; Fig. 8C). Frequency distribution analyses indicated that complex-shaped AS were larger than macular AS in all cortical regions (KS; P<0.01). The relatively low number of SS precluded us from performing a robust statistical analysis.

### Size of axospinous and axodendritic synapses

To determine whether the synaptic size was related to the postsynaptic targets, we analyzed the SAS area of both AS and SS.

Analyses of the mean SAS area and the frequency distribution showed that AS established on spine heads were significantly larger than those established on dendritic shafts without spines in BA17, BA3b and BA4, and also larger than those on dendritic shafts with spines in BA17 and BA4 (KW, P<0.05; KS, P<0.05; Fig. 8D; Supplementary Fig. 4; Supplementary Table 7). In BA17, larger AS were more frequently found on dendritic shafts with spines than on shafts without spines (KS; P<0.05; Supplementary Fig. 4; Supplementary Table 7). Since the number of SS on spine heads was relatively low (ranging from 3 to 8 in the analyzed cortical regions), this statistical analysis was discarded as it would not have been sufficiently robust.

Further comparisons of all previously examined cortical regions were carried out (BA24, vBA38, dBA38, and BA21; Cano-Astorga et al., 2023). It was found that the AS established on spines were larger than AS on dendritic shafts in all cortical regions except in BA3b and BA24. BA4, vBA38, and BA21 displayed the largest difference in the AS size between spines and shafts, whereas BA3b and BA24 had similar SAS areas of AS in spines and shafts (Fig. 8D). The number of SS on complete spine heads (ranging from 3 to 14 in the cortical regions analyzed) was not large enough to perform a robust statistical analysis.

Since the synaptic size correlates with release probability, synaptic strength, efficacy, and plasticity (see Chindemi et al., 2022 and references therein), to better understand differences between cortical regions, we determined a new ratio to estimate the total AS SAS area per region. This “SAS area/volume ratio” was calculated by dividing the sum of all the SAS areas from AS in each cortical region by their analyzed volume (corresponding to the inclusion volume of the CF). This ratio revealed that BA4 had lower values than BA24, vBA38, dBA38, and BA21 (KW; P<0.05), whereas BA4 showed no differences compared to BA17 and BA3b (Supplementary Table 8; Supplementary Fig. 5A). Additionally, the values were assigned to two groups: BA17, BA3b, and BA4 — and BA24, vBA38, dBA38, and BA21. Comparing the “SAS area/volume ratio” of AS, significant differences were found between the two groups (MW; P<0.0001), with lower values in the former (BA17, BA3b, and BA4).

## Interindividual variability

Variability between individuals was examined by calculating the coefficient of variation (CV) of each synaptic parameter in every cortical region (except BA17).

We observed the highest variability in synaptic density between individuals in BA3b (CV=25%; Table 2). By contrast, BA21 exhibited remarkably homogeneous synaptic density across individuals (CV=8%; Table 2). The proportion of AS was homogenous among individuals (Table 2). Exploring the mean of the AS SAS area revealed the highest interindividual variability in BA24 (CV=23%; Table 2), while BA4 yielded remarkably similar values across individuals (CV=5%; Table 2). The estimated “SAS area/volume ratio” indicated that, out of all of the cortical regions studied, BA21 had the lowest variability (CV=3%; Table 2). Regarding the synaptic shape, the percentage of macular AS was remarkably homogenous among individuals (Table 2), with CVs ranging from 0.28 to 7.08%. Finally, the percentage of AS established on spines varied by about 15% between individuals in both BA3b and BA4, while the lowest variation was observed in BA24 (2.64%; Table 2).

**Table 2.**
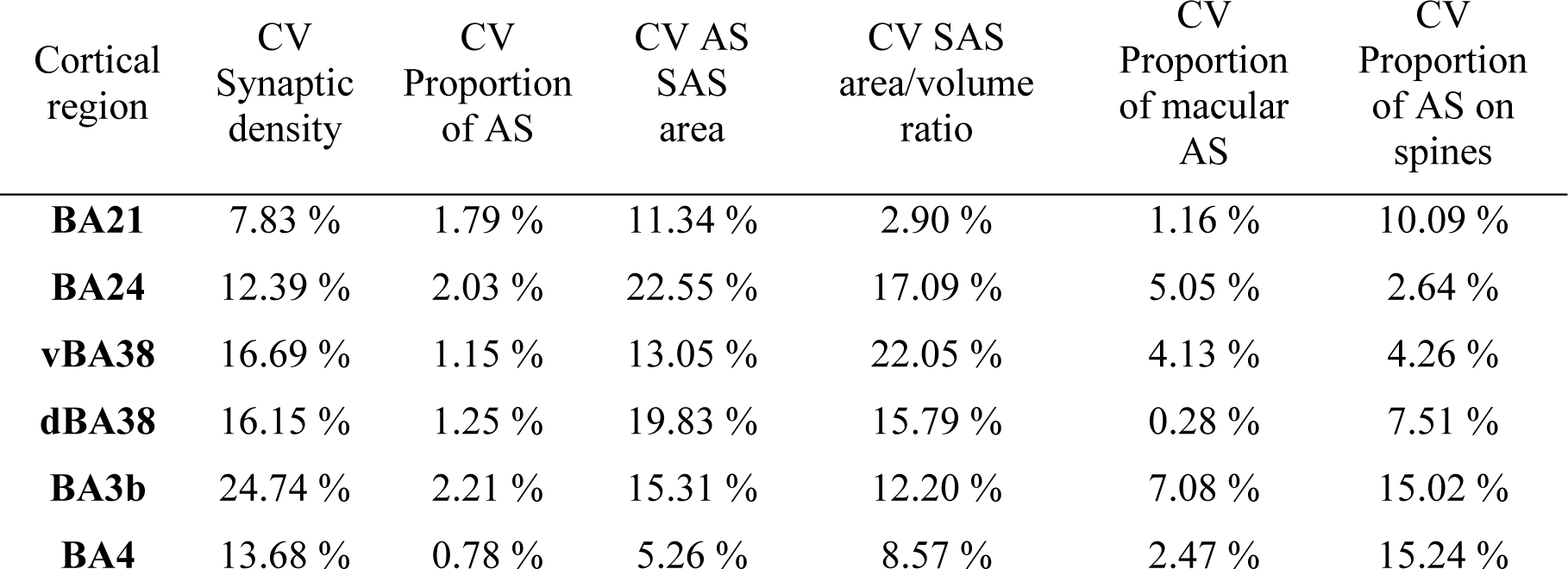
CV between cases of different synaptic parameters (columns) in each cortical region (rows). CV is expressed as percentage. AS: asymmetric synapses; BA: Brodmann area; CV: coefficient of variation; d: dorsal; SAS: synaptic apposition surface; v: ventral.

| Cortical region | CV Synaptic density | CV Proportion of AS | CV AS SAS area | CV SAS area/volume ratio | CV Proportion of macular AS | CV Proportion of AS on spines |
| --- | --- | --- | --- | --- | --- | --- |
| **BA21** | 7.83 % | 1.79 % | 11.34 % | 2.90 % | 1.16 % | 10.09 % |
| **BA24** | 12.39 % | 2.03 % | 22.55 % | 17.09 % | 5.05 % | 2.64 % |
| **vBA38** | 16.69 % | 1.15 % | 13.05 % | 22.05 % | 4.13 % | 4.26 % |
| **dBA38** | 16.15 % | 1.25 % | 19.83 % | 15.79 % | 0.28 % | 7.51 % |
| **BA3b** | 24.74 % | 2.21 % | 15.31 % | 12.20 % | 7.08 % | 15.02 % |
| **BA4** | 13.68 % | 0.78 % | 5.26 % | 8.57 % | 2.47 % | 15.24 % |

Therefore, in general, we observed that interindividual variability is associated with certain cortical regions and specific synaptic characteristics.

## Discussion

The present results provide a large quantitative ultrastructural dataset of synapses in layer III of the human primary visual, somatosensory and motor cortices using 3D EM. This study also compares the present results with our previous studies focusing on different cortical regions (BA24, vBA38, dBA38 and BA21), performed in the same individuals.

The main finding is that there are certain synaptic characteristics —such as number of synapses per volume, postsynaptic target distribution, and the synaptic size of AS—which seem to be specific to cortical regions. However, other characteristics are common to all analyzed regions, including: i) AS:SS ratio; ii) the random spatial distribution of the synapses in the neuropil; iii) SAS area of AS being larger than that of SS; iv) the size of SS; v) the macular shape of most synapses, and their small size compared to complex-shaped synapses; vi) most synapses being AS established on spines, followed by AS on dendritic shafts and SS on dendritic shafts; and vii) most dendritic spine heads receiving a single AS. To sum up, some synaptic traits are specific to certain cortical regions, while other synaptic features are common across them.

## Region-specific characteristics

### Synaptic number and density

Variations in the Vv-neuropil may imply significant differences in the total number of synapses, as suggested by DeFelipe et al. (1999). For instance, BA17 and BA3b had comparable synaptic densities to those observed in BA24, BA21, and vBA38, ranging from 0.44 to 0.52 synapses/μm^3^. However, the Vv-neuropil in the former group of cortical areas was approximately 75%, which was lower than in the latter group, which had an approximate Vv-neuropil of 90%. Hence, when considering both synaptic density and Vv-neuropil, it is evident that layer III of BA17 and BA3b are likely to contain a lower number of synapses compared to BA24, BA21, and vBA38. Additionally, given that BA4 has the lowest synaptic density and a low Vv-neuropil, this area is expected to have fewer synapses than BA17, BA3b, BA24, BA21 and vBA38. Moreover, all of these regions are anticipated to have a lower number of synapses than dBA38, which has the highest synaptic density and a high Vv-neuropil. Thus, understanding the overall connectivity of these cortical regions requires consideration of the significance of volume fraction variations in conjunction with the synaptic density and the total volume of the layer.

Given that the majority of connections in the cerebral cortex are formed by point-to-point chemical synapses (DeFelipe, 2015), assessing the synaptic density in a particular region is crucial for understanding both connectivity and functionality. In the present study, we observed heterogeneity in the mean synaptic density of the neuropil in layer III across the cortical areas analyzed. Previous studies employing the same techniques have demonstrated variations in synaptic density in the neuropil of other brain regions, including layer II of the transentorhinal cortex (Domínguez-Álvaro et al., 2018) and layers II and III of the entorhinal cortex (Domínguez-Álvaro et al., 2021a). Notably, these synaptic densities were found to be comparable to those observed in BA24, vBA38, BA21, BA3b and BA17. Nevertheless, the deep and superficial stratum pyramidale of the CA1 field of the hippocampus —from the same individuals as those examined in the current study— exhibited a higher synaptic density (ranging from 0.67 to 0.99 synapses/μm^3^; Montero-Crespo et al., 2020). This range aligns more closely with the synaptic density observed in dBA38. No other 3D EM study has reported a mean synaptic density as low as that discovered in BA4 (0.28 synapses/μm^3^) in human brain samples. Given the identical tissue processing and analysis methods, any similarities or differences are likely attributable to specific characteristics of the brain region and layer being analyzed. Furthermore, the lowest mean distance to the nearest synapse is associated with dBA38, characterized by the highest synaptic density. By contrast, BA4, with the lowest synaptic density, has the highest mean distance to the nearest synapse.

The current findings on synaptic density, along with the Vv-neuropil, are in line with previous reports of higher spine density and pyramidal cell complexity in layer III from BA38, BA21, BA24, and BA17 (Benavides-Piccione et al., 2013, 2021, 2023). Unfortunately, data on the morphology of layer III pyramidal neurons in the primary somatosensory (BA3b) and motor cortex (BA4) of the human cerebral cortex are not available. Concerning non-human primates, studies have indicated that spine density and the branching complexity of pyramidal neurons are lower in the primary sensory-motor cortex compared to other associative cortices, such as the anterior cingulate and temporal cortices (Elston et al., 2005a, 2005b). Further research is necessary to explore potential correlations of dendritic spine density and synaptic density across different human cortical areas.

### Postsynaptic Target Distribution

In the present study, the concurrent analysis of the synaptic type and postsynaptic target revealed that the proportions of axospinous AS were approximately 73–77% in BA24, vBA38, dBA38, BA21, and BA3b. However, in BA17 and BA4, a lower proportion was observed — 68% and 62%, respectively. Comparisons of the proportions of AS established on spines in other cortical regions in the same autopsy cases, using the same FIB/SEM technology, revealed heterogeneous proportions. For example, in BA24, vBA38, dBA38, BA21 and BA3b, proportions of axospinous AS are similar to those obtained in layer II of the human transentorhinal cortex (75%; Domínguez-Álvaro et al., 2021a), as are the axospinous AS proportions in the stratum oriens, deep stratum pyramidale, and stratum radiatum of the CA1 hippocampal field (77–81%; Montero-Crespo et al., 2020). However, in layer II of the human entorhinal cortex, the proportion of AS established on spines was reported to be 60% (Domínguez-Álvaro et al., 2021a), similar to in BA17 and BA4. Furthermore, the proportion of axospinous AS was lower in layer III of the human entorhinal cortex (56%; Domínguez-Álvaro et al., 2021a) and in the stratum lacunosum-moleculare of the CA1 hippocampal field (57%; Montero-Crespo et al., 2020). Additionally, the superficial stratum pyramidale of the CA1 hippocampal field had the highest proportion of axospinous AS reported to date (88%; Montero Crespo et al., 2020).

Therefore, these variations in the proportion of AS on spines and dendritic shafts signify an additional microanatomical specialization in the examined cortical regions (and layers).

These differences in synaptic organization may have significant functional implications for the processing of information in these cortical regions. For instance, studies, including Cornejo et al. (2022) and references therein, have indicated that the membrane potential of the postsynaptic neuron is differentially modulated depending on whether a synapse is established on a dendritic spine or a dendritic shaft. Consequently, variations in the proportions of these targets have functional significance.

### Synaptic Size

It has been proposed that synaptic size correlates with release probability, synaptic strength, efficacy, and plasticity (see Chindemi et al., 2022 and references therein). Several methods have traditionally been used to estimate the size of synaptic junctions, making it difficult to compare between different studies (reviewed in Cano-Astorga et al., 2021). Analysis of the SAS area, conducted on a substantial number of axospinous AS (approximately 33,000 synaptic junctions) in both the present and previous studies (Domínguez-Álvaro et al., 2018; 2021a; Montero-Crespo et al., 2020), constitutes a robust 3D morphological dataset of synapses. This analysis reveals significant differences between cortical regions. For instance, the SAS area of AS in BA17 and BA3b is smaller (approximately 100,000 nm^2^) than in BA24, BA21 and vBA38, which share similar values to those obtained in layer II of the transentorhinal cortex (Domínguez-Álvaro et al., 2018), and layers II and III of the entorhinal cortex (Domínguez-Álvaro et al., 2021a). dBA38 had an SAS area of approximately 80,000 nm^2^, notably lower than that of BA24, BA21, and vBA38, and comparable to values obtained in all layers of the CA1 field of the hippocampus (Montero-Crespo et al., 2020). BA4 had a mean SAS area of around 130,000 nm^2^, which was the largest among the cortical regions analyzed to date.

It has been demonstrated that synaptic size correlates with the number of receptors in the PSD; larger PSDs have more receptors (as reviewed in Lüscher et al., 2000; Lüscher and Malenka, 2012; Toni et al., 2001; Magee and Grienberger, 2020; Sumi and Harada, 2020). Therefore, the aforementioned differences in the size of PSDs and synaptic density between cortical areas are generally in line with previous research indicating significant variations among cortical areas (including primary sensory, motor, and multimodal association areas) regarding the density (measured as fmol mg/protein) of various transmitter systems, such as glutamate, GABA, acetylcholine, dopamine, noradrenaline, and serotonin (Zilles and Palomero-Gallagher, 2017, and references therein).

Nevertheless, it has been reported that some receptors can be found both synaptically and extra-synaptically (Palomero-Gallagher and Zilles, 2019). Therefore, caution should be exercised when considering potential correlations between receptor densities, number of synapses and/or synaptic size.

Further analysis of the AS based on their postsynaptic target revealed that those established on dendritic spine heads were larger than those established on dendritic shafts (with and without spines) in BA4, BA17, vBA38, dBA38, and BA21. This observation is in line with previous findings reported in layer II of the transentorhinal cortex (Domínguez-Álvaro et al., 2019) and layer II and III of the entorhinal cortex (Domínguez-Álvaro et al., 2021a). However, in BA3b and BA24, the size of AS established on spine heads and those on dendritic shafts (with and without spines) was similar, consistent with findings in the CA1 field of the hippocampus (Montero-Crespo et al., 2020). This observation reinforces the notion of regional synaptic specialization, suggesting potential functional implications in the information processing across distinct cortical regions.

## Common synaptic characteristics

### Proportion of AS:SS and spatial synaptic distribution

It is well established that the cortical neuropil has a larger proportion of AS than SS regardless of the cortical region and species. Transmission electron microscopy studies have shown that this proportion varies between 80 and 95% for AS and 20–5% for SS (reviewed in DeFelipe et al., 2002; Bourne and Harris, 2011). In the present study, the AS:SS ratio varies between 92 and 94% for AS and 8-6% for SS, which is similar to the reported data in other human cortical regions with FIB/SEM (Domínguez-Álvaro et al., 2018, 2021a; Montero-Crespo et al., 2020; Cano-Astorga et al., 2021, 2023). Hence, the AS:SS ratio, as revealed by 3D electron microscopy, generally indicates a higher proportion of AS and a lower proportion of SS compared to previous findings using transmission electron microscopy.

Interpretation of the significance of the consistent AS:SS ratio across cortical regions proves challenging, given the variations in cytoarchitecture, neurochemistry, connectivity, and functional characteristics among different brain regions. These regions host a diverse array of neurochemical and functional types of pyramidal and GABAergic neurons and it has been shown that there are differences in the number of GABAergic and glutamatergic inputs in several neuronal types (e.g., DeFelipe and Fariñas, 1992; Freund and Buzsáki, 1996; DeFelipe, 1997; Somogyi et al., 1998; Schubert et al., 2007; Markram et al., 2015; Tremblay et al., 2016; Hu et al., 2018; Sohal and Rubenstein, 2019; Chini et al., 2022). Thus, examining the synaptic inputs of each specific cell type becomes essential for determining differences in the AS:SS ratio, even if the overall AS:SS proportion in the neuropil remains constant. Finally, it is worth noting that our focus has been on the synaptic organization of the neuropil, excluding perisomatic innervation. The study of perisomatic innervation is also crucial for a comprehensive understanding of the synaptic organization within cortical circuits, as highlighted in a recent study by Ostos et al. (2023).

Regarding the spatial organization of synapses, it was observed that synapses were randomly distributed in the neuropil across most of the samples of BA17, BA3b, and BA4. This pattern is consistent with findings in the transentorhinal, entorhinal, temporopolar, and anterior cingulate cortices, as well as in CA1 field of the hippocampus (Domínguez-Álvaro et al., 2018, 2021a; Montero-Crespo et al., 2020; Cano-Astorga et al., 2023). As emphasized in a previous study (Merchán-Pérez et al., 2014), the random distribution of synapses should not be misconstrued as indicating random connections between neurons, as our data reflect a heterogeneous population of synapses. Spatial randomness does not necessarily mean non-specific connections. For example, diverse neuronal types have preferences for establishing synapses with different types of neurons and/or different parts of these neurons (such as different segments of the dendritic tree, spines, dendritic shafts, soma, or axon initial segment), showing specific connectivity patterns. Consequently, our conclusion that synapses are randomly distributed in space should be interpreted in the context of the entire heterogeneous population of synapses.

### AS are larger than SS

While most studies generally estimate the size of synapses, particularly AS, accurate data on the size of SS are scarce. In the current study, we observed that AS were larger than SS in layer III from BA17, BA3b and BA4, consistent with previous findings in other human cortical regions, including layer III of BA24, BA38, and BA21 (Cano-Astorga et al., 2023), layer II of the transentorhinal cortex (Domínguez-Álvaro et al., 2018), layers II and III of the entorhinal cortex (Domínguez-Álvaro et al., 2021a), and in all layers of the CA1 field of the hippocampus (Montero-Crespo et al., 2020).

Applying the same techniques to other mammalian species, AS were found to be larger than SS in all layers of the somatosensory cortex of the Etruscan Shrew (Alonso-Nanclares et al., 2022). However, it has been demonstrated that SS are larger than AS in certain layers of the CA1 field of the mouse hippocampus (Santuy et al., 2020), in certain layers of the mouse somatosensory cortex (Turégano-López, 2022), and in all layers of the juvenile rat somatosensory cortex (Santuy et al., 2018a). Hence, conducting additional studies across diverse brain regions, layers, and species is necessary to investigate the presence of a regional and/or species-specific pattern in the mammalian cerebral cortex.

### Synaptic size of SS

In addition, the analysis of the SAS area in SS revealed no differences between the cortical regions analyzed in the present study — and this was also the case when comparing with other human cortical regions analyzed in previous studies (Domínguez-Álvaro et al., 2018, 2021a; Montero-Crespo et al., 2020; Cano-Astorga et al., 2023). Hence, SS in the neuropil appear to constitute a genuinely homogeneous population of synapses. Moreover, SAS area values of SS were found to be comparable to those obtained in various cortical regions of other mammals, such as the primary somatosensory cortex of the Etruscan shrew (60,378 nm^2^; Alonso-Nanclares et al., 2022), the primary somatosensory cortex of the mouse (67,281 nm^2^; Turégano-López, 2022), and the stratum lacunosum-moleculare, stratum radiatum and stratum oriens of the hippocampal field of the mouse (54,100 nm^2^; Santuy et al., 2020).

Conducting further studies in the neuropil of other cortical areas, layers and species will help determine whether the size of the SS can be regarded as a regional and/or species-specific characteristic of the mammalian cerebral cortex.

### Most synapses are macular-shaped, and they are the smallest

We found that the majority of synapses had a macular shape (79–87%) and were the smallest synapses. These findings are in line with previous reports in other brain regions and species (Geinisman et al., 1987; Jones et al., 1991; Neuman et al., 2016; Hsu et al., 2017; Calì et al., 2018; Santuy et al., 2018a; Domínguez-Álvaro et al., 2019, 2021a; Montero-Crespo et al., 2020; Cano-Astorga et al., 2021, 2023; Turégano-López, 2022; Alonso-Nanclares et al., 2023). It has been reported that complex-shaped synapses possess more AMPA and NMDA receptors than macular synapses, characterizing them as a potentially ’powerful’ population associated with more enduring memory-related functionality than macular synapses (Geinisman et al., 1987, 1991, 1992a, 1992b, 1993; Lüscher et al., 2000; Toni et al., 2001; Ganeshina et al., 2004a, 2004b; Spruston 2008). Conversely, the smaller area of macular synapses may play a crucial role in synaptic plasticity (Kharazia and Weinberg, 1999). Thus, from the functional point of view, determining the shape of synapses can offer valuable insights. While further studies may yield slightly different results, currently, the prevalence of approximately 70–90% macular small synapses and 10–30% large complex-shaped synapses could be considered as another specific characteristic of the mammalian cerebral cortex.

### Most synapses are axospinous AS followed by axodendritic AS and then axodendritic SS

A clear preference of glutamatergic axons (forming AS) targeting spines, and GABAergic axons (forming SS) targeting dendritic shafts, was observed in the present study, consistent with numerous electron microscopy studies across various cortical regions and species (reviewed in DeFelipe et al., 2002). However, a frequent misunderstanding regarding this characteristic is that it suggests that synapses on dendritic shafts are predominantly SS. In reality, quantitative analyses of synapses in the neuropil reveal that the majority of synapses are AS established on spines, followed by AS on dendritic shafts, and then SS on dendritic shafts (Beaulieu et al., 1992; Peters et al., 2008; Hsu et al., 2017; Calì et al., 2018; Santuy et al., 2018b; Yakoubi et al., 2019; Montero-Crespo et al., 2020, 2021; Domínguez-Álvaro et al., 2019, 2021a, 2021b; Cano-Astorga et al., 2021, 2023; Schmuhl-Giesen et al., 2022; Turégano-López, 2022). Thus, the prevalence of axospinous AS, followed by axodendritic AS and axodendritic SS, can be considered an additional characteristic among the general rules governing human synapses.

### Most spines receive a single AS

Regarding the number of synapses per dendritic spine, our findings indicate that the majority of spines receive a single AS across all cortical regions analyzed. The proportion of spines receiving a single AS in BA3b (95.2%) and in BA17 (92.2%) was comparable to values obtained in layer III of BA24, vBA38, dBA38 and BA21 (range: 93.4% to 96.2%; Cano-Astorga et al., 2023) and layer II of the transentorhinal cortex (94.5%; Domínguez-Álvaro et al., 2019). By contrast, the proportion observed in BA4 (89.6%) was comparable to that found in layers II and III of the entorhinal cortex (90.3% and 89.6%, respectively; Domínguez-Álvaro et al., 2021a). Additionally, in the CA1 field of the hippocampus, the percentage was even higher (97.7%; Montero-Crespo et al., 2020). Hence, the observation that most spines receive a single AS constitutes another fundamental characteristic of human synapses.

The functional significance of having single or multiple synapses on the same spine is yet to be fully understood. The formation of multi-synaptic spines has been linked to synaptic potentiation and memory processes (Giese et al., 2015; McLeod et al., 2020). In the mouse neocortex, it has been suggested that spines receiving one AS and one SS are electrically more stable than spines with a single AS (Villa et al., 2016). Given that most spines are single-innervated spines, it is conceivable that multisynaptic spines represent a distinct postsynaptic element. For instance, in the cerebral cortex of monkeys and humans, it has been demonstrated that in a specific type of GABAergic cell, known as the double bouquet cell, approximately 38–48% of the synapses (SS) are established on spines that also receive an additional AS (Somogyi and Cowey, 1981; DeFelipe et al., 1989, 1990; De Lima and Morrison, 1989; del Rio and DeFelipe, 1995; Lukacs et al., 2022). This represents a remarkably high percentage of SS formed on spines, especially considering that dual-innervated spines constitute only 5–10% of the total population of synapses in the neuropil.

## Relationship between the synaptic organization and the structural/functional macroscopic connectivity

Functional and structural studies investigating macroscopic connectivity in the human cerebral cortex have suggested that high-order associative cortex, such as BA24, BA38 and BA21, exhibited greater connectivity compared to primary cortex (including BA17, BA3b and BA4) (Sporns et al., 2005; Van Essen et al., 2013; Paquola et al., 2020). The calculated “SAS area/volume ratio” also supports this notion, indicating a higher synaptic surface in the high-order associative regions compared to the primary cortex (Supplementary Fig. 5). Given the strong correlation between SAS area and synaptic strength, the “SAS area/volume ratio” implies increased synaptic activity in associative cortex (BA24, vBA38. dBA38, and BA21) compared to primary cortex (BA17, BA3b and BA4). However, when we consider both the density and size of synapses, we observe regions with high synaptic densities but small sizes, while other regions have lower synaptic densities but larger synaptic sizes. This significant variation in synaptic connectivity distinguishes one region from another. In addition, the proportion of macular and complex-shaped synapses, along with the distribution and size of synapses on spines and dendritic shafts, plays a fundamental role with regard to the dynamism and computation of the neural circuits. Therefore, it is crucial to consider all morphological characteristics of synapses to provide accurate insights into the connectivity of the cerebral cortex. Hence, an integrative multiscale study at micro-, meso-, and macroscopic levels would be necessary to obtain a more comprehensive understanding of the synaptic organization.

## Interindividual variability of the human cerebral cortex

A number of studies highlighted the interindividual variability in the structural and functional organization of the human brain (Jacobs and Scheibel, 1993; Benavides-Piccione et al., 2005, 2013, 2021, 2023; Benavides-Piccione and DeFelipe, 2007; Alonso-Nanclares et al., 2008; Blázquez-Llorca et al., 2010; Fernández-González et al., 2016; Peng et al., 2019; Montero-Crespo et al., 2020; Domínguez-Álvaro et al., 2021a; Cano-Astorga et al., 2021, 2023). In the present study, we observed that interindividual variability exists at the level of synaptic characteristics, but this variability is associated with certain cortical regions and specific synaptic characteristics. Some regions exhibited more variability in synaptic density (e.g., dBA38 and BA3b), while others were more variable with regard to other synaptic characteristics such as the proportion of macular and complex-shaped synapses (e.g., BA24), or the distribution of postsynaptic targets (e.g., BA21 and BA3b). Thus, although there are synaptic characteristics that are common among different regions of the human cerebral cortex, which may be considered as general rules of synaptic organization, interindividual variability should also be taken into account depending on the cortical region and specific synaptic characteristics examined.

## Conclusions

The present study provides a large quantitative ultrastructural dataset of synapses in layer III of the regions analyzed. The results highlight the presence of some regional-specific synaptic characteristics, while other synaptic features are common across regions. These findings may contribute to a better understanding of the organization of the human cerebral cortex and help decipher its connectivity more accurately.

## Author contributions

N. Cano-Astorga: Conceptualization, Formal analysis, Investigation, Methodology, Visualization, Writing - original draft, Writing - review & editing; S. Plaza-Alonso: Investigation, Validation, Writing - review & editing; J. DeFelipe: Conceptualization, Funding acquisition, Resources, Supervision, Writing - review & editing; L. Alonso- Nanclares: Conceptualization, Data curation, Methodology, Project Administration, Supervision, Validation, Writing - review & editing.

## Classification

Research reports; direct submission

## Acknowledgements

We would like to thank Nick Guthrie for his excellent editorial assistance.

## Funding

Grant PID2021-127924NB-I00 funded by MCIN/AEI/10.13039/501100011033 (J.D.)

CSIC Interdisciplinary Thematic Platform - Cajal Blue Brain (J.D.)

Research Fellowships funded by MCIN/AEI/10.13039/501100011033 for N.C.-A. (PRE2019-089228) and S.P.-A. (FPU19/00007).

## Ethics approval

Brain tissue samples were obtained following the guidelines and approval of the Institutional Ethical Committee at the School of Medicine, University of Castilla-La Mancha (Albacete, Spain).

## Availability of data and materials

Most data are available in the main text and the Supplementary Data. Some of the datasets used and analyzed during the current study are published in the EBRAINS Knowledge Graph:

Domínguez-Álvaro M, Montero M, Alonso-Nanclares L, Blazquez-Llorca L, Rodríguez J-R, DeFelipe J. (2020). Densities and 3D distributions of synapses using FIB/SEM imaging in the human neocortex (Temporal cortex, T2). Human Brain Project Neuroinformatics Platform. (DOI: 10.25493/5F04-N97).

Alonso-Nanclares L., Cano-Astorga N., Plaza-Alonso S., DeFelipe J. (2022). “3D ultrastructural study of synapses using FIB/SEM in the Human Cortex (Brodmann areas 24 and 38)”. Human Brain Project Neuroinformatics Platform. (DOI: 10.25493/B3V0- 4D8).

## Abbreviation list

3D: three-dimensional
AMPA: α-amino-3-hydroxy-5-methyl-4-isoxazolepropionic acid
ANOVA: analysis of variance
AS: asymmetric synapses
CA1: Cornu Ammonis 1
CF: counting frame
CSR: Complete Spatial Randomness
CV: coefficient of variation
d: dorsal
FIB/SEM: focused ion beam/scanning electron microscopy
KS: Kolmogorov-Smirnov
KW: Kruskal-Wallis
MW: Mann-Whitney
NMDA: N-methyl-D-aspartate
PB: phosphate buffer
PSD: postsynaptic density
SAS: synaptic apposition surface
SE: standard error of the mean
SD: standard deviation
SS: symmetric synapses
TEM: transmission electron microscopy
v: ventral
Vv: volume fraction
χ^2^: chi-square

## Supplementary Data

**Supplementary Table 1.** Summary of the stack details obtained from the multiple sampling of BA17, BA3b, BA4, BA24, BA38v, BA38d, and BA21 in the human autopsy cases AB2, AB3 and AB7. Data between parentheses indicate the sum of the number of slices and the volume examined per stack. The results from the analyses of BA24, BA38v, BA38d, and BA21 were published previously in Cano-Astorga et al., (2023).

| Cortical region | Cases | No. of stacks of images | Distance from pial surface ($\mu\text{m}$; range) | Total volume of analyzed neuropil ($\mu\text{m}^3$) | No. of slices per stack (range; mean) | Volume examined per stack ($\mu\text{m}^3$; range; mean) |
| --- | --- | --- | --- | --- | --- | --- |
| **BA17** | AB7 | 4 | 600 | 1,763 | 261-305; 280 | 411-480; 441 |
| **B3b** | AB2, AB3, AB7 | 9 | 600-700 | 4,034 | 252-306; 285 | 396-481; 448 |
| **BA4** | AB2, AB3, AB7 | 9 | 600-900 | 4,119 | 281-318; 287 | 445-500; 458 |
| **BA24** | AB2, AB3, AB7 | 9 | 600-1100 | 4,289 | 240-330; 273 | 377-519; 429 |
| **vBA38** | AB2, AB3, AB7 | 9 | 700-800 | 4,399 | 254-314; 280 | 400-494; 440 |
| **dBA38** | AB2, AB3, AB7 | 9 | 600-900 | 4,336 | 261-305; 276 | 411-480; 434 |
| **BA21** | AB2, AB3, AB7 | 7 | 400-600 | 3,896 | 260-317; 287 | 409-499; 451 |
| *All* | *AB2, AB3, AB7* | *56* | *400-1100* | *26,836* | *(15,765)* | *(19,844)* |

**Supplementary Table 2.**
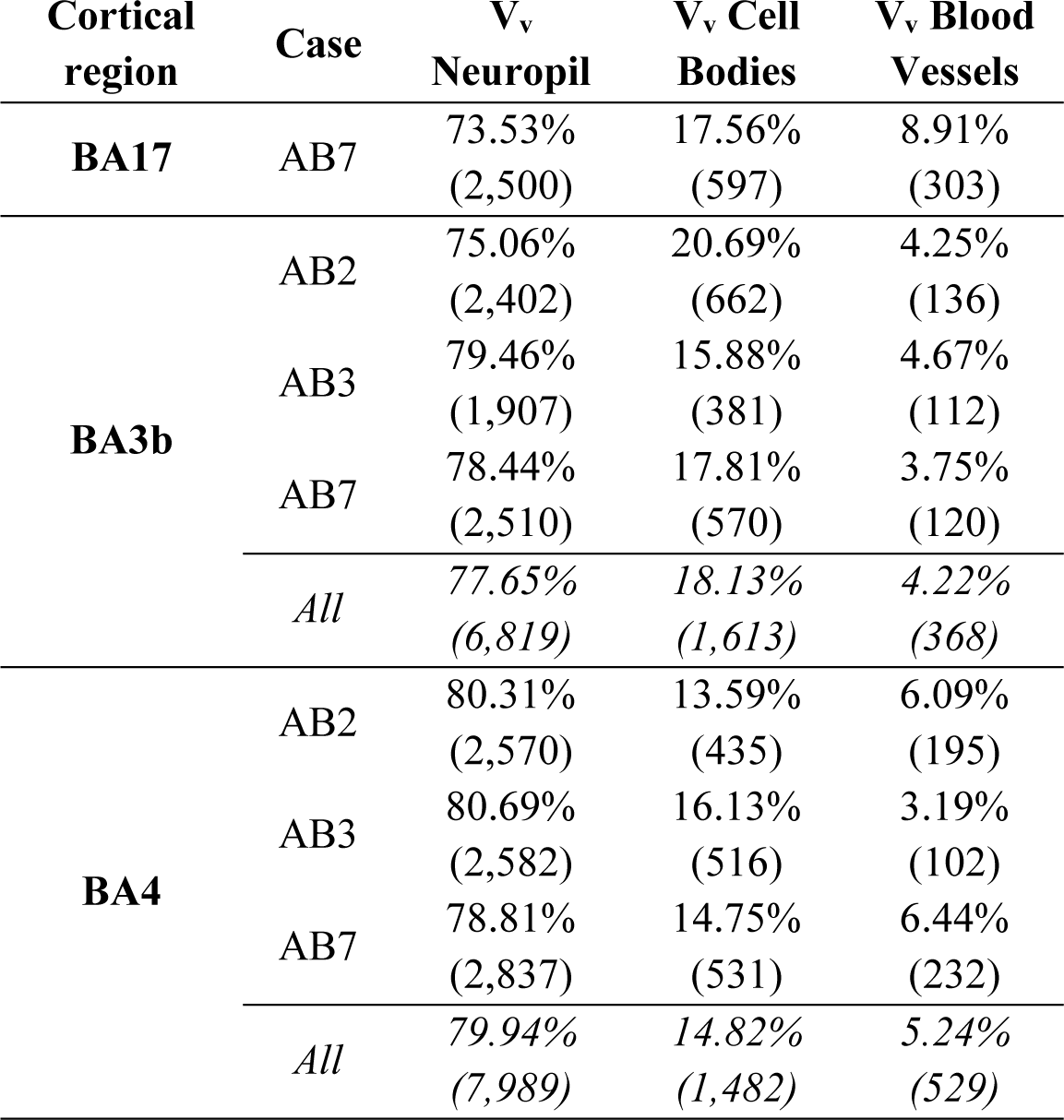
Volume fraction (V_v_) occupied by different cortical elements in BA17, BA3b, and BA4 per case. The absolute numbers below the percentages of volume fractions indicate the number of times that a grid point was counted in each category (neuropil, cell bodies and blood vessels). BA: Brodmann area.

**Supplementary Table 3.**
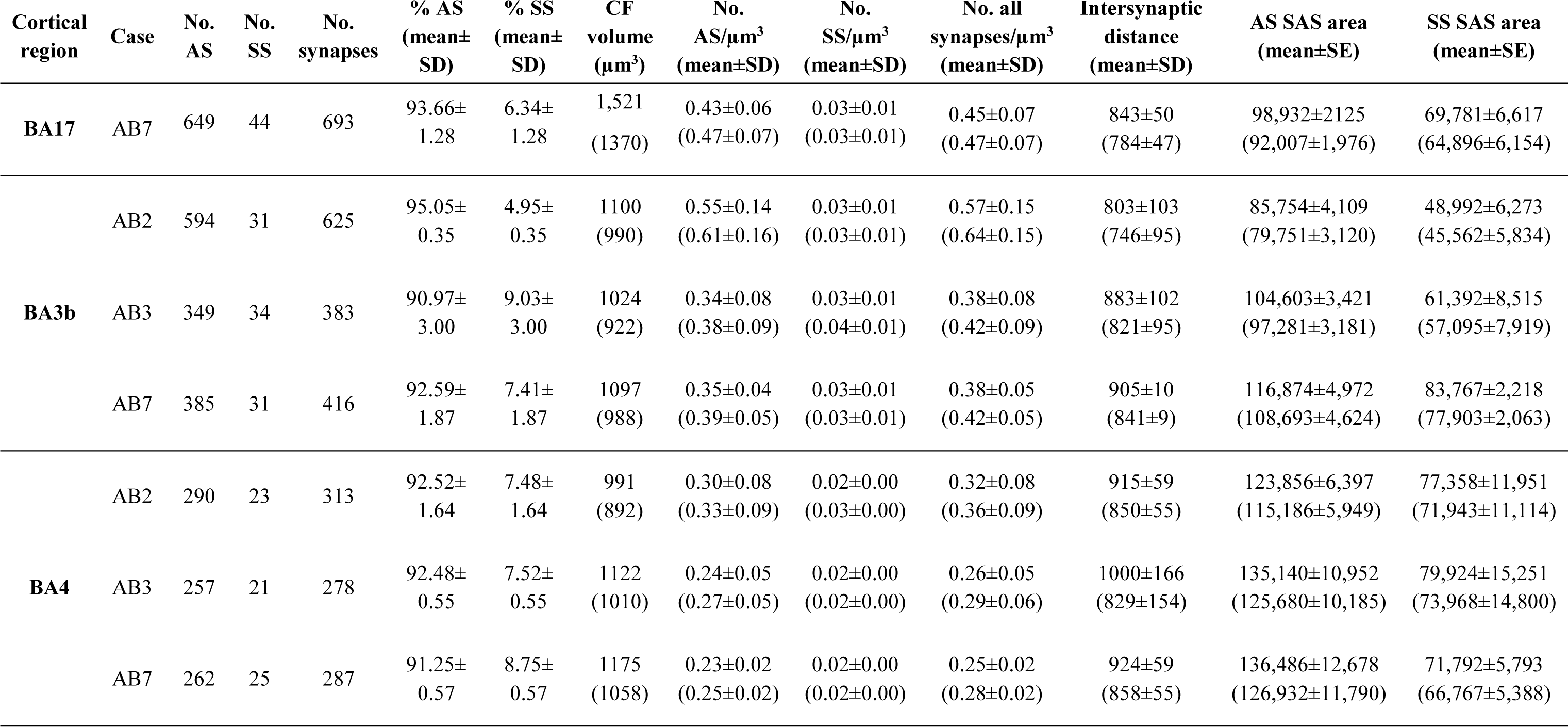
Accumulated data obtained from the ultrastructural analysis of neuropil from layer III of BA17, BA3b and BA4 per case. Data in parentheses have not been corrected for shrinkage. AS: asymmetric synapses; BA: Brodmann area; CF: counting frame; SAS: synaptic apposition surface; SD: standard deviation; SE: standard error of the mean; SS: symmetric synapses.

**Supplementary Table 4.** Proportions of the different synaptic shapes from layer III of BA17, BA3b and BA4 per case. Data are given as percentages (absolute numbers of synapses studied are given in parentheses). AS: asymmetric synapses; BA: Brodmann area; SS: symmetric synapses.

| Cortical region | Case | Type of synapse | Macular synapses | Perforated synapses | Horseshoe-shaped synapses | Fragmented synapses | Total synapses |
| --- | --- | --- | --- | --- | --- | --- | --- |
| **BA17** | **AB7** | AS | 87.2%<br>(566) | 8.6%<br>(56) | 3.1%<br>(20) | 1.1%<br>(7) | 100%<br>(649) |
|  |  | SS | 79.5%<br>(35) | 6.8%<br>(3) | 11.4%<br>(5) | 2.3%<br>(1) | 100%<br>(44) |
| **BA3b** | **AB2** | AS | 82.8%<br>(492) | 12.8%<br>(76) | 4.2%<br>(25) | 0.2%<br>(1) | 100%<br>(594) |
|  |  | SS | 90.3%<br>(28) | 6.5%<br>(2) | 0.0%<br>(0) | 3.2%<br>(1) | 100%<br>(31) |
|  | **AB3** | AS | 79.7%<br>(278) | 12.9%<br>(45) | 6.6%<br>(23) | 0.9%<br>(3) | 100%<br>(349) |
|  |  | SS | 88.2%<br>(30) | 0.0%<br>(0) | 5.9%<br>(2) | 5.9%<br>(2) | 100%<br>(34) |
|  | **AB7** | AS | 88.1%<br>(339) | 7.3%<br>(28) | 4.4%<br>(17) | 0.3%<br>(1) | 100%<br>(385) |
|  |  | SS | 84.6%<br>(22) | 11.5%<br>(3) | 3.8%<br>(1) | 0.0%<br>(0) | 100%<br>(26) |
|  | **All** | AS | 83.5%<br>(1109) | 11.2%<br>(149) | 4.9%<br>(65) | 0.4%<br>(5) | 100%<br>(1328) |
|  |  | SS | 87.9%<br>(80) | 5.5%<br>(5) | 3.3%<br>(3) | 3.3%<br>(3) | 100%<br>(91) |
| **BA4** | **AB2** | AS | 80.3%<br>(233) | 15.5%<br>(45) | 3.8%<br>(11) | 0.3%<br>(1) | 100%<br>(290) |
|  |  | SS | 82.6%<br>(19) | 8.7%<br>(2) | 8.7%<br>(2) | 0.0%<br>(0) | 100%<br>(23) |
|  | **AB3** | AS | 76.7%<br>(197) | 19.1%<br>(49) | 2.7%<br>(7) | 1.6%<br>(4) | 100%<br>(257) |
|  |  | SS | 90.5%<br>(19) | 9.5%<br>(2) | 0.0%<br>(0) | 0.0%<br>(0) | 100%<br>(21) |
|  | **AB7** | AS | 79.8%<br>(209) | 13.7%<br>(36) | 3.4%<br>(9) | 3.1%<br>(8) | 100%<br>(262) |
|  |  | SS | 92.0%<br>(23) | 0.0%<br>(0) | 0.0%<br>(0) | 8.%<br>(2) | 100%<br>(25) |
|  | **All** | AS | 79.0%<br>(639) | 16.1%<br>(130) | 3.3%<br>(27) | 1.6%<br>(13) | 100%<br>(809) |
|  |  | SS | 88.4%<br>(61) | 5.8%<br>(4) | 2.9%<br>(2) | 2.9%<br>(2) | 100%<br>(61) |

**Supplementary Table 5.**
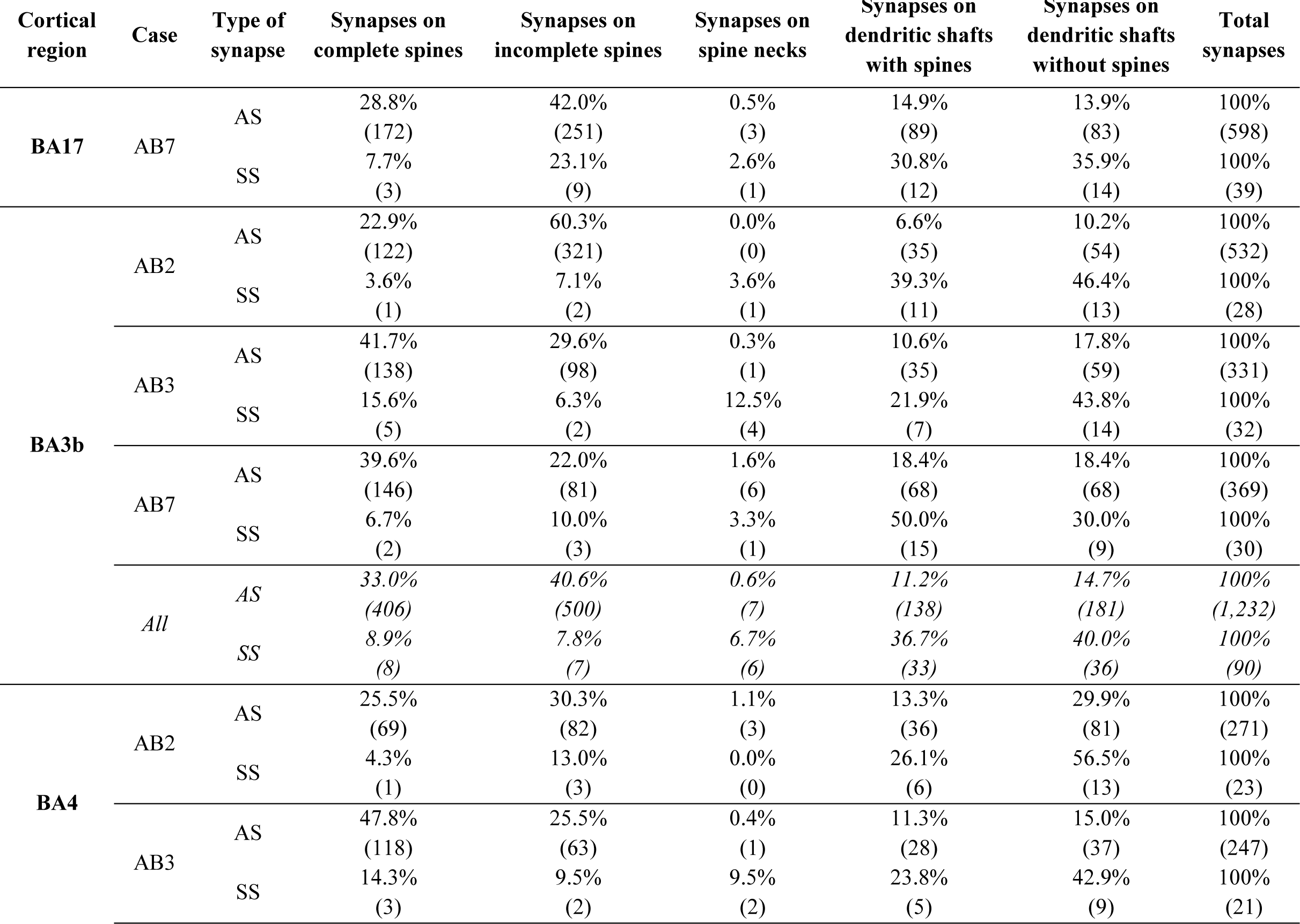
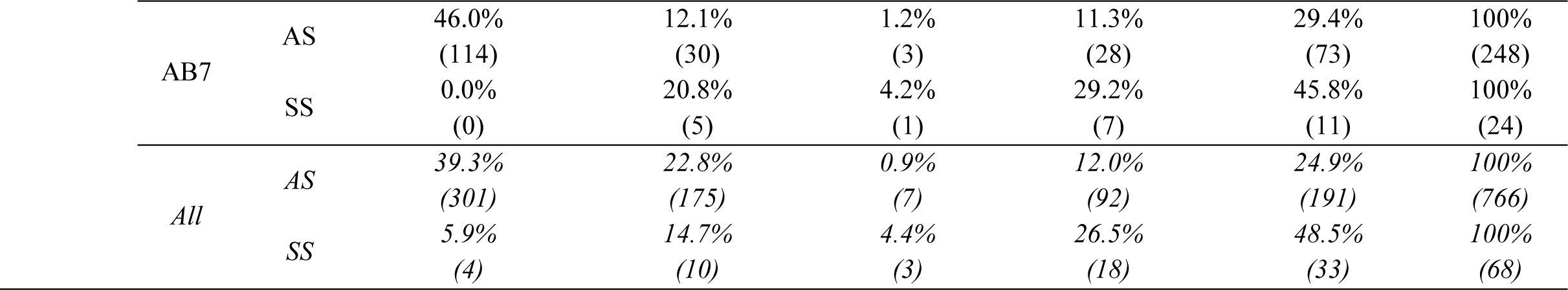
Distribution of AS and SS on spines and dendritic shafts in layer IIIA of BA17, BA3b and BA4 per case. Synapses on spines have been sub-divided into those that are established on spine heads and those established on spine necks. Data are given as percentages with the absolute number of synapses studied in parentheses. AS: asymmetric synapses; BA: Brodmann area; SS: symmetric synapses.

**Supplementary Table 6.** SAS area (nm^2^) of macular, perforated, horseshoe and fragmented AS and SS in layer III of BA17, BA3b and BA4. SAS areas of the macular AS were significantly smaller than perforated and horseshoe in all regions (KW, p<0.001), and smaller than fragmented AS in BA17 and BA4 (KW; P<0.001). Data in parentheses are not corrected for shrinkage. AS: asymmetric synapses; BA: Brodmann area; NA: not applicable; SAS: synaptic apposition surface; SE: standard error of the mean; SS: symmetric synapses.

| Cortical region | Type of synapses | Morphological category | SAS area (mean ± SE) |
| --- | --- | --- | --- |
| BA17 | AS | Macular | 89,537 ± 1,913 (83,270 ± 1,779) |
| | | Perforated | 164,927 ± 8,543 (153,382 ± 7,945) |
| | | Horseshoe | 143,034 ± 10,096 (133,022 ± 9,389) |
| | | Fragmented | 204,219 ± 35,018 (189,924 ± 32,567) |
| | SS | Macular | 61,854 ± 6,050 (57,524 ± 5,627) |
| | | Perforated | 52,602 ± 7,892 (48,920 ± 7,340) |
| | | Horseshoe | 126,615 ± 26,256 (117,752 ± 24,418) |
| | | Fragmented | 179,345 ± NA (166,791 ± NA) |
| BA3b | AS | Macular | 88,620 ± 1,495 (82,417 ± 1,391) |
| | | Perforated | 154,375 ± 5,402 (143,569 ± 5,024) |
| | | Horseshoe | 150,018 ± 6,921 (139,517 ± 6,437) |
| | | Fragmented | 147,435 ± 23,038 (137,114 ± 21,425) |
| | SS | Macular | 55,182 ± 4,291 (51,320 ± 3,991) |
| | | Perforated | 109,813 ± 31,294 (102,127 ± 29,103) |
| | | Horseshoe | 92,623 ± 53,062 (86,140 ± 49,348) |
| | | Fragmented | 55,960 ± 11,885 (52,043 ± 11,053) |
| BA4 | AS | Macular | 104,855 ± 2,523 (97,515 ± 2,347) |
| | | Perforated | 231,747 ± 9,211 (215,525 ± 8,567) |
| | | Horseshoe | 222,503 ± 10,999 (206,928 ± 10,229) |
| | | Fragmented | 210,460 ± 16,281 (195,728 ± 15,142) |
| | SS | Macular | 71,481 ± 5,686 (66,477 ± 5,288) |
| | | Perforated | 129,441 ± 24,997 (120,380 ± 23,247) |
| | | Horseshoe | 53,716 ± 24,345 (49,956 ± 22,641) |
| | | Fragmented | 108,929 ± 17,455 (101,304 ± 16,233) |

**Supplementary Table 7.** SAS area (nm^2^) of AS and SS established on complete spine heads, incomplete spine heads, spine necks, dendritic shafts with spines and dendritic shafts without spines in layer III of BA17, BA3b, and BA4. The area of the AS established on complete spine heads was significantly larger than the area of the AS established on dendritic shafts (with spines and without spines) in all regions except BA3b. Data in parentheses are not corrected for shrinkage. AS: asymmetric synapses; BA: Brodmann area; NA: not applicable; SAS: synaptic apposition surface; SE: standard error of the mean; SS: symmetric synapses.

| Cortical region | Type of synapses | Postsynaptic target | SAS area (mean ± SE) |
| --- | --- | --- | --- |
| BA17 | AS | Complete spine heads | 118,044 ± 4,307 (109,781 ± 4,006) |
| | | Incomplete spine heads | 95,448 ± 3,314 (88,766 ± 2,914) |
| | | Spine necks | 107,337 ± 24,270 (99,823 ± 22,571) |
| | | Shafts with spines | 99,880 ± 6,878 (92,889 ± 6,397) |
| | | Shafts without spines | 82,231 ± 4,855 (76,475 ± 4,516) |
| | SS | Complete spine heads | 52,745 ± 27,695 (49,053 ± 25,756) |
| | | Incomplete spine heads | 59,251 ± 10,777 (55,103 ± 10,023) |
| | | Spine necks | 19,313 ± NA (17,961 ± NA) |
| | | Shafts with spines | 79,605 ± 13,543 (74,033 ± 12,595) |
| | | Shafts without spines | 73,225 ± 13,599 (68,099 ± 12,647) |
| BA3b | AS | Complete spine heads | 106,239 ± 2,708 (98,802 ± 2,518) |
| | | Incomplete spine heads | 93,270 ± 2,387 (86,741 ± 2,220) |
| | | Spine necks | 65,593 ± 16,944 (64,722 ± 15,758) |
| | | Shafts with spines | 110,189 ± 5,281 (102,476 ± 4,911) |
| | | Shafts without spines | 94,924 ± 5,147 (88,279 ± 4,786) |
| | SS | Complete spine heads | 53,509 ± 7,202 (49,763 ± 6,698) |
| | | Incomplete spine heads | 46,352 ± 12,300 (43,107 ± 11,439) |
| | | Spine necks | 58,577 ± 23,109 (54,477 ± 21,491) |
| | | Shafts with spines | 75,505 ± 9,629 (70,220 ± 8,955) |
| | | Shafts without spines | 62,117 ± 7,769 (57,769 ± 7,255) |
| BA4 | AS | Complete spine heads | 148,150 ± 4,776 (137,779 ± 4,442) |
| | | Incomplete spine heads | 137,746 ± 6,645 (128,104 ± 6,180) |
| | | Spine necks | 79,754 ± 21,712 (74,171 ± 20,193) |
| | | Shafts with spines | 111,152 ± 8,574 (103,371 ± 7,974) |
| | | Shafts without spines | 114,632 ± 6,829 (106,608 ± 6,351) |
| | SS | Complete spine heads | 65,091 ± 8,382 (60,535 ± 7,795) |
| | | Incomplete spine heads | 69,975 ± 10,521 (65,077 ± 9,785) |
| | | Spine necks | 46,175 ± 21,986 (42,943 ± 20,447) |
| | | Shafts with spines | 59,810 ± 9,617 (55,623 ± 8,944) |
| | | Shafts without spines | 85,129 ± 9,102 (79,170 ± 8,465) |

**Supplementary Table 8.**
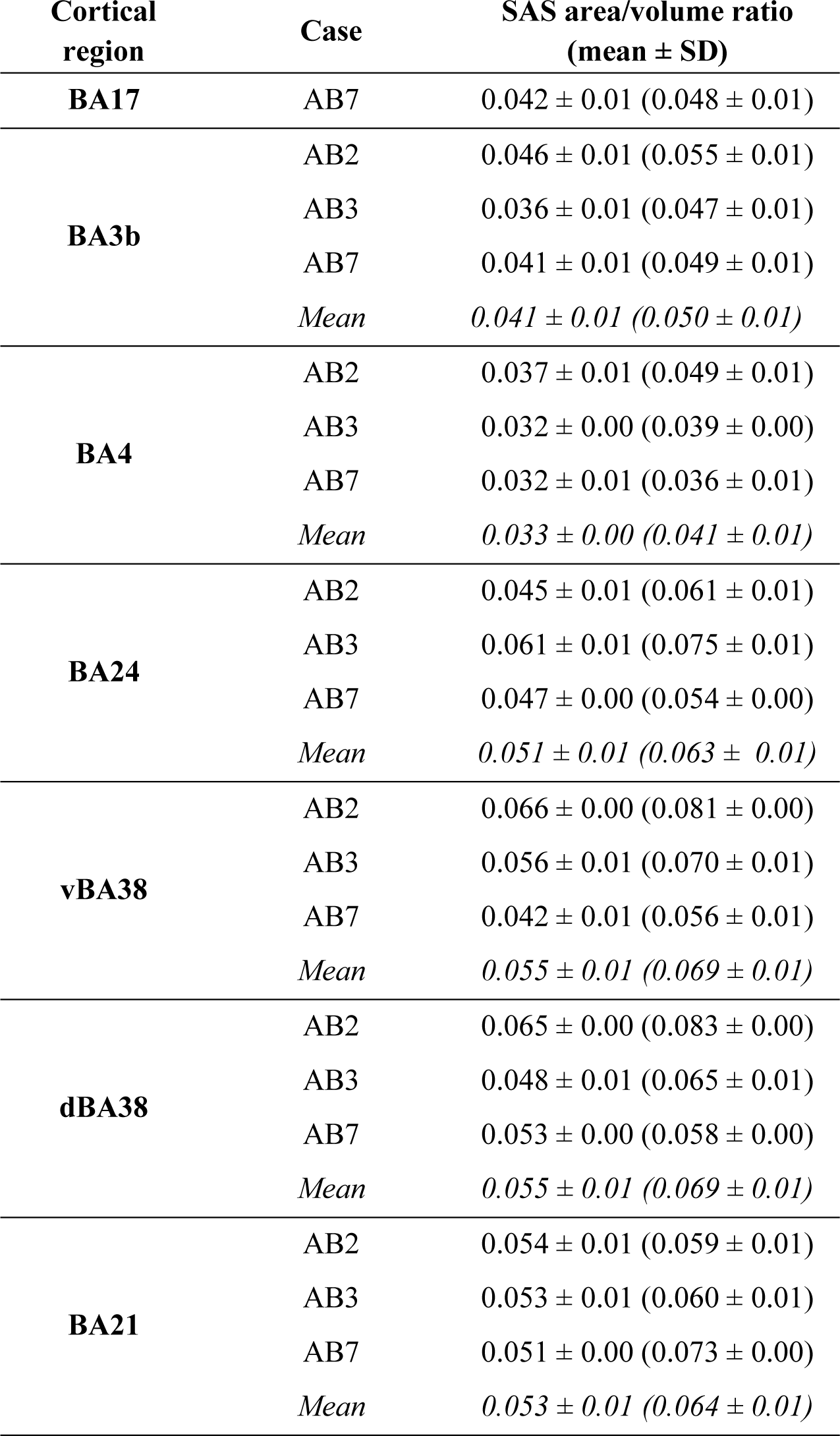
SAS area/volume ratio of BA17, BA3b, BA4, BA24, vBA38, dBA38, and BA21. Data are given per case and the mean are given in italics. Data in parentheses have not been corrected for shrinkage. AS: asymmetric synapse; BA: Brodmann area; d: dorsal; SAS: synaptic apposition surface; Vv: volume fraction; v: ventral.

**Supplementary Figure 1.**
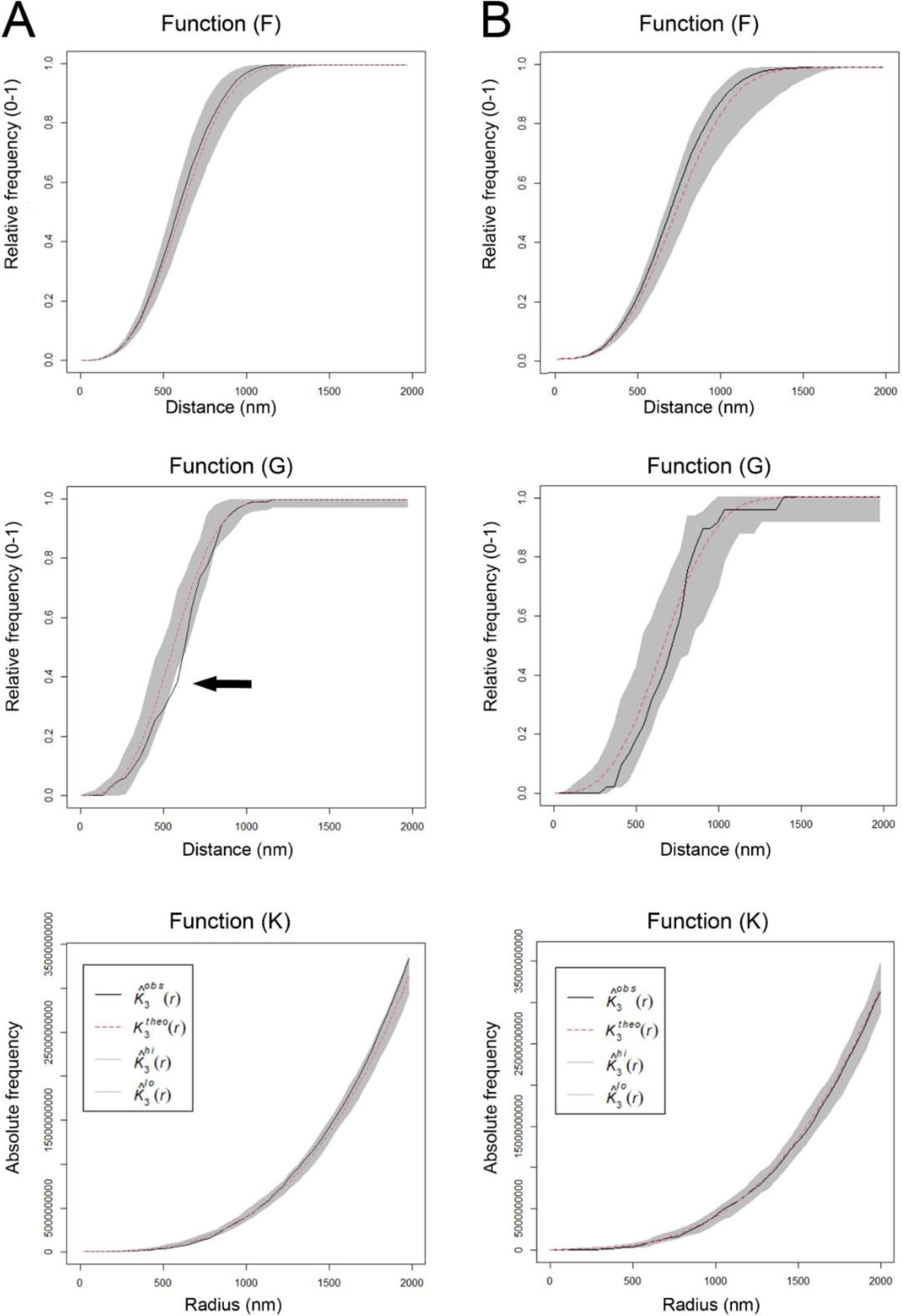
Analysis of the 3D synaptic spatial distribution in layer III of BA17, BA3b, and BA4. Red dashed traces correspond to a theoretical homogeneous Poisson process for each function (F, G, K). The black continuous traces correspond to the experimentally observed function in the sample. The shaded areas represent the envelopes of values calculated from a set of 99 simulations. (A) Example of stacks (2 out of 22) showing a slight tendency toward a distributed pattern in one of the functions indicated by the arrow — in this case, the G function. (B) Example of stacks (20 out of 22) fitting a Poisson function.

**Supplementary Figure 2.**
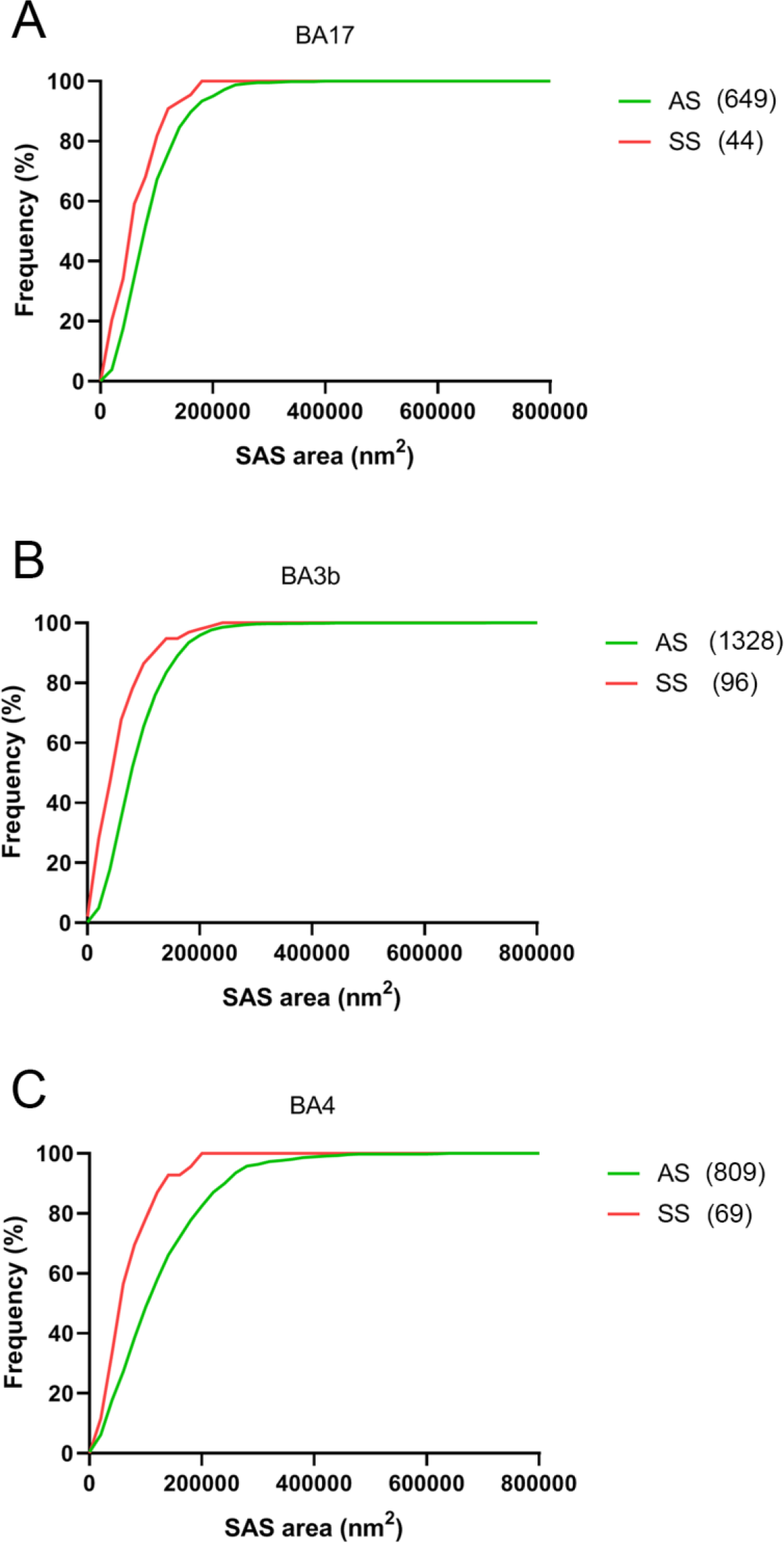
Cumulative frequency distribution of SAS area of AS and SS. (A–G) Small SS (red) were more frequent than small AS (green) in all regions (KS P<0.001). AS: asymmetric synapses; BA: Brodmann area; SAS: synaptic apposition surface; SS: symmetric synapses.

**Supplementary Figure 3.**
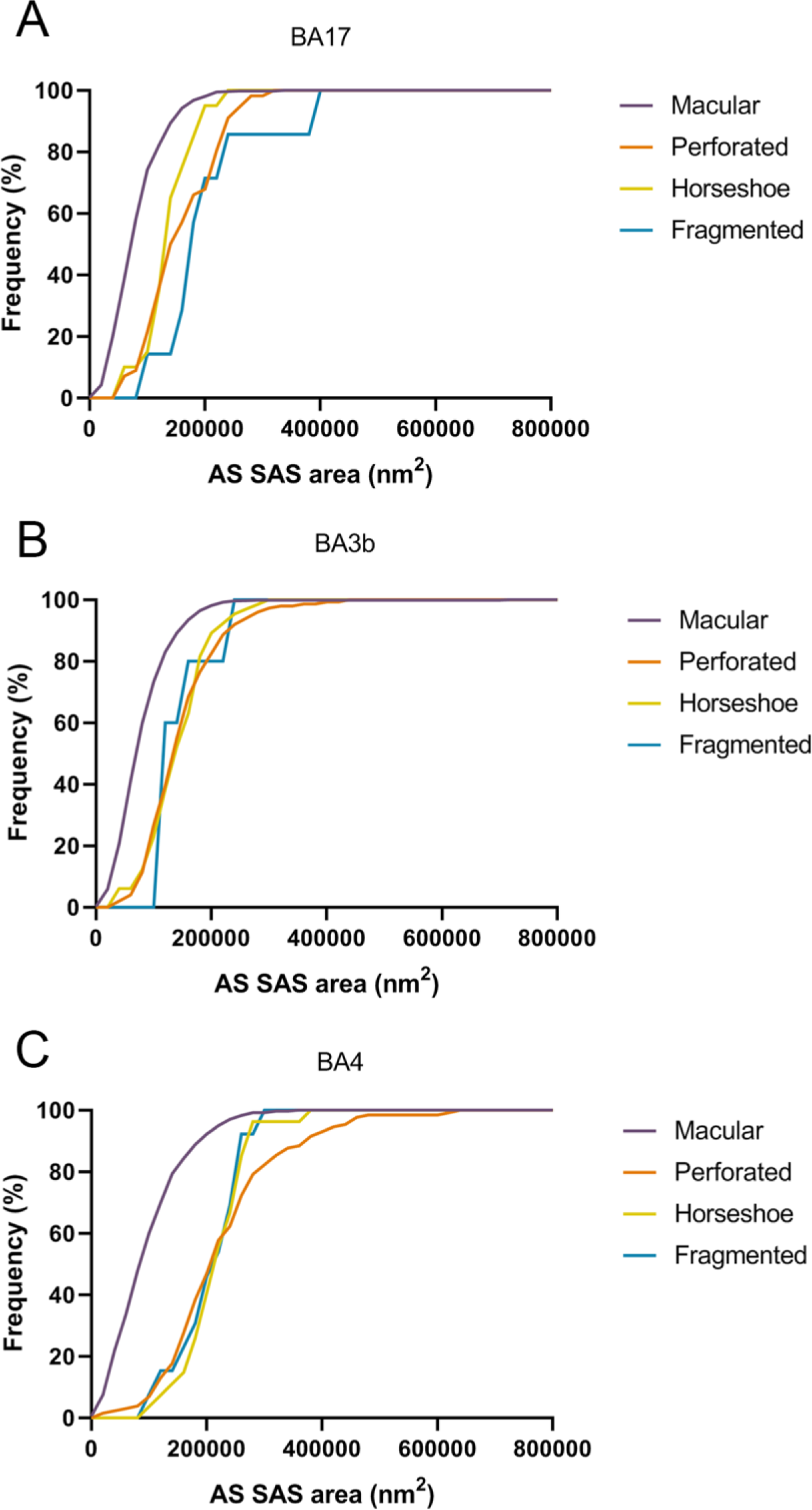
Frequency histograms of the SAS area of AS per synaptic shape from layer III of BA17 (A), BA3b (B). Smaller macular AS were the most frequent shape in all regions (KS, p<0.0001). AS: asymmetric synapses; BA: Brodmann area; SAS: synaptic apposition surface..

**Supplementary Figure 4.**
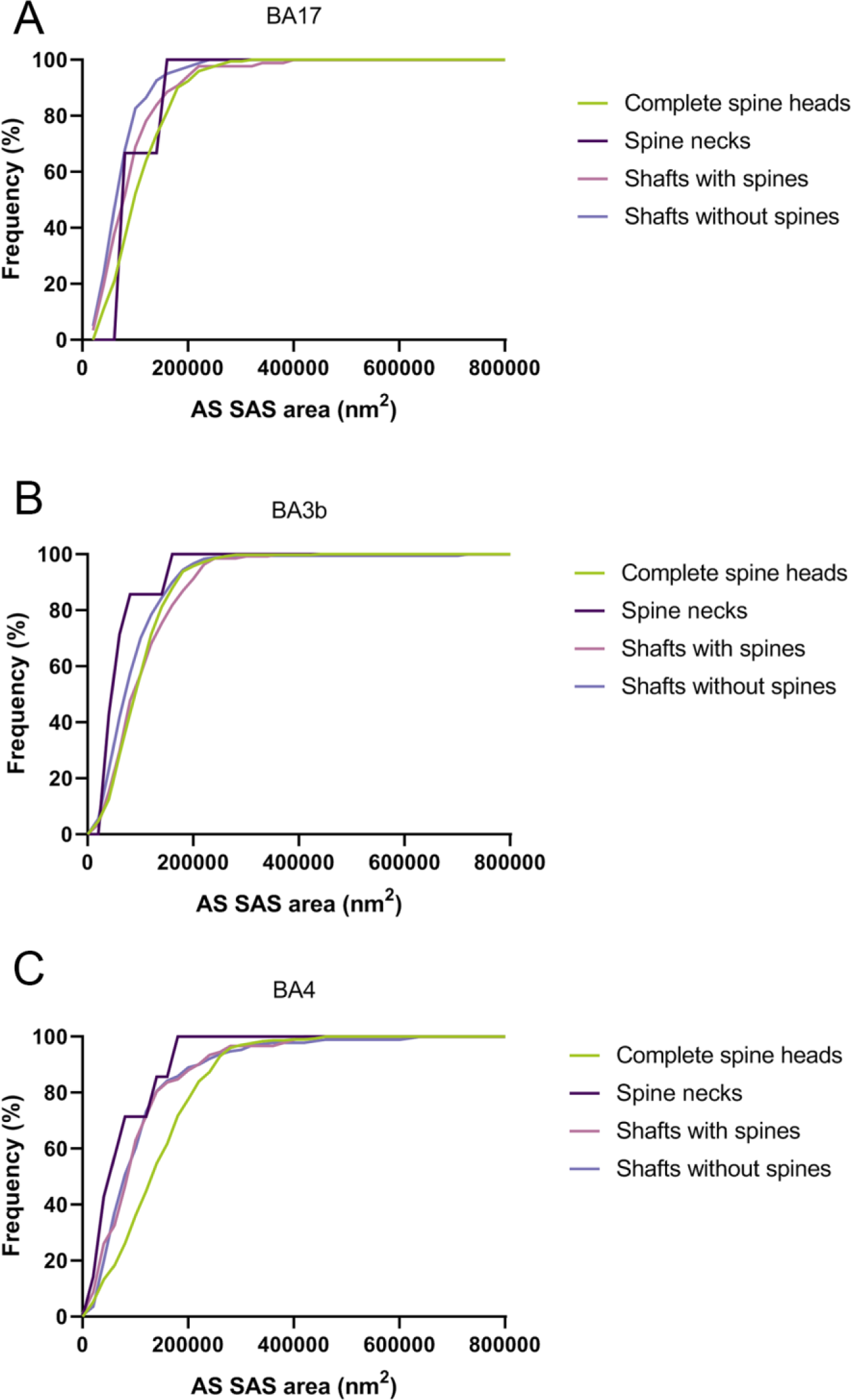
Frequency histograms of the SAS area of Asymmetric Synapses established on complete spines, spine necks, shafts with spines and shafts without spines from layer III neuropil of BA17 (A), BA3b (B) and BA4 (C) samples. The larger SAS areas were significantly more frequent in AS on complete spine heads than in the case of AS on dendritic shafts without spines in all cortical regions (KS: P<0.01), AS on dendritic shafts with spines in BA17 and BA4 (KS; P<0.01). In addition, the larger SAS areas were more frequently found in AS on dendritic shafts with spines than in AS on dendritic shafts without spines in BA17 (KS; P<0.01). BA: Brodmann area; SAS: synaptic apposition surface.

**Supplementary Figure 5.**
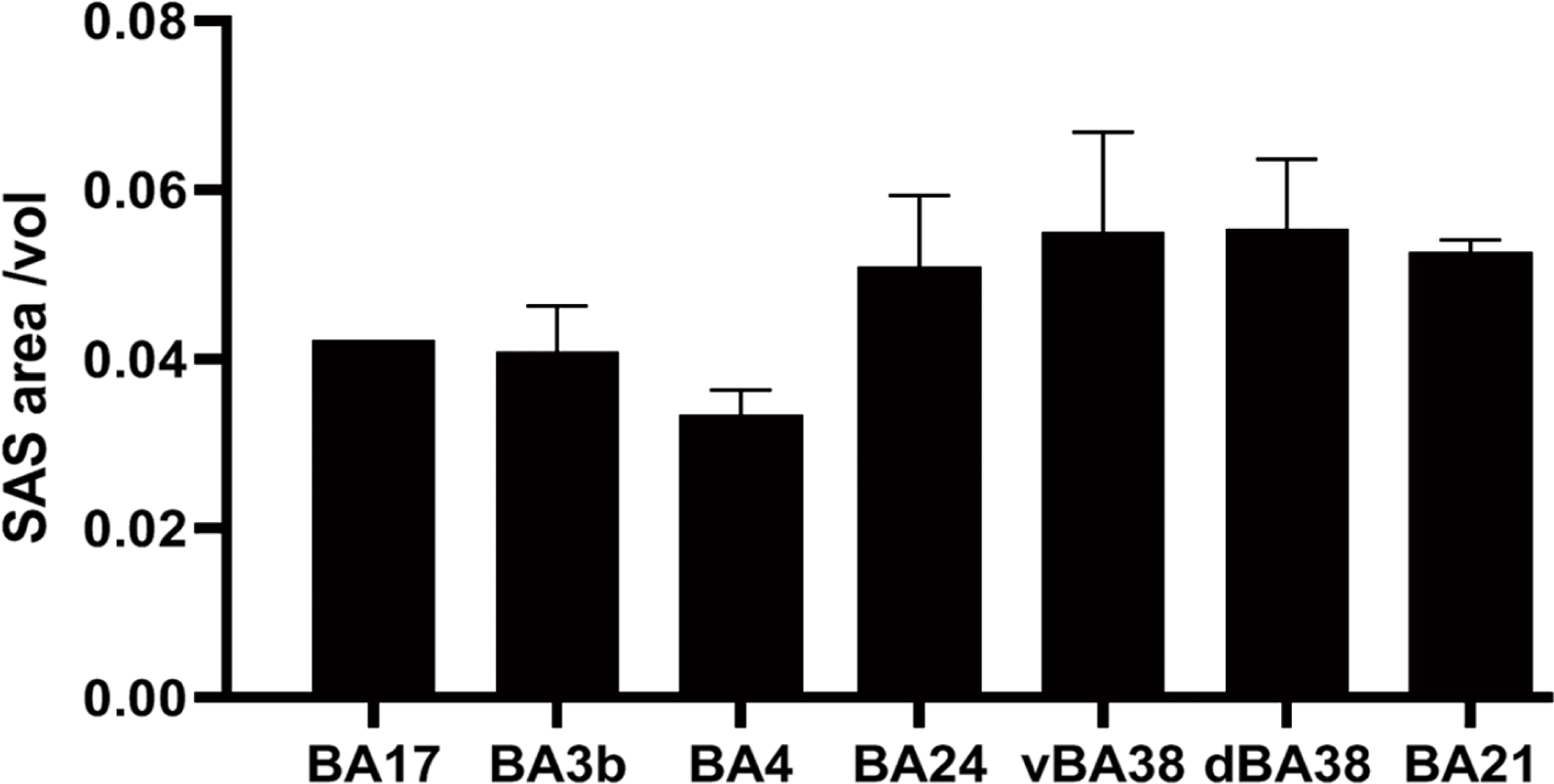
Plot of the SAS area/volume ratio in BA17, BA3b, BA4, BA24, vBA38, dBA38, and BA21 from three individuals (mean ± SD). BA17 data come from the analysis of one individual. BA17, BA3b, and BA4 showed lower values than BA24, vBA38, dBA38, and BA21, and significant differences were found (KW; P<0.05) when comparing BA4 with BA24, vBA38, dBA38, and BA21. AS: asymmetric synapses; BA: Brodmann area; d: dorsal; SAS: synaptic apposition surface; SD: standard deviation; SS: symmetric synapses; v: ventral.

